# Reconciling competing models on the roles of condensates and soluble complexes in transcription factor function

**DOI:** 10.1101/2024.11.21.624739

**Authors:** Anne Bremer, Walter H. Lang, Ryan P. Kempen, Kumari Sweta, Aaron B. Taylor, Madeleine B. Borgia, Aseem Z. Ansari, Tanja Mittag

## Abstract

Phase separation explains the exquisite spatial and temporal regulation of many biological processes, but the role of transcription factor–mediated condensates in gene regulation is contentious, requiring head-to-head comparison of competing models. Here, we focused on the prototypical yeast transcription factor Gcn4 and assessed two models for gene transcription activation, i.e., mediated via soluble complexes or transcriptional condensates. Both models rely on the ability of transcription factors and coactivators to engage in multivalent interactions. Unexpectedly, we found that propensity to form homotypic Gcn4 condensates does not correlate well with transcriptional activity. Contrary to prevailing models, binding to DNA suppresses Gcn4 phase separation. Notably, the ability of Gcn4 to form soluble complexes with coactivator subunit Med15 closely mirrored the propensity to recruit Med15 into condensates, indicating that these properties are intertwined and cautioning against interpretation of mutational data without head-to-head comparisons. However, Gcn4 variants with the highest affinity for Med15 do not function as well as expected and instead have activities that reflect their abilities to phase separate with Med15. These variants therefore indeed form cellular condensates, and those attenuate activity. Our results show that transcription factors can function as soluble complexes as well as condensates, reconciling two seemingly opposing models, and have implications for other phase-separating systems.

**Highlights:**

- Homotypic phase separation propensities of Gcn4 variants do not predict *in vivo* activities well.
- DNA binding leads to solubilization of GCN4 condensates, which is reversed by Med15 association.
- The abilities to co-phase separate and form soluble complexes with Med15 are highly intertwined.
- Variants with high affinities for Med15 form condensates that attenuate function.

## Main

Transcription requires the coordinated recruitment of transcription factors (TFs), coactivators and the transcriptional machinery to enhancers and promoters ^1^. Recent work proposes that the coordination of these events is achieved via DNA-scaffolded phase separation, in which TFs and the transcriptional machinery engage enhancers and promoters through the formation of biomolecular condensates ^2-9^. According to these models, multiple DNA sites serve as scaffolds that facilitate the formation of TF condensates ^2,10-12^. Phase separation is reported to coordinate transcriptional activation ^3,7^, elongation ^13^, termination ^14^, splicing ^15^, as well as transcriptional repression ^16-20^. While this paradigm provided new directions to investigate the assembly of transcriptional machinery and transcription-related pathogenic processes, there is still intense debate on whether phase separation is necessary, sufficient, and consequential in modulating transcriptional output ^3-5,15,21-27^.

The proposal that phase separation plays a role in transcriptional regulation was initially based on multiple properties shared by super-enhancers and phase-separating systems ^2^. First, the clustering of multiple enhancer elements in super-enhancers mediates multivalent interactions that can scaffold the formation of condensates. Second, super-enhancers are characterized by all-or-nothing, switch-like responses that turn transcription on/off upon small changes in TF concentration; the extreme cooperativity of phase separation lends itself to generating such all-or-nothing responses if the dense phase is considerably more active than the system in the absence of phase separation ^28,29^. Indeed, DNA constructs with multiple binding sites for the pioneer factor Oct4 or the repressive MeCP2 protein scaffold their phase separation ^10,18^. Third, coactivators such as the Mediator complex undergo phase separation with pioneer factors Oct4, Sox2, and others ^3^. Fourth, specific Mediator subunits are sufficient to enhance TF phase separation ^4^. And fifth, mutant TFs with diminished ability to drive phase separation show markedly reduced transcriptional output, and this correlation was taken as evidence that phase separation is causative and essential for transcription activation ^3,18,23,30,31^.

The concept that phase separation mediates transcriptional control was extended to promoters that are not under the control of super-enhancers. For instance, the YAP/TAZ TFs that drive developmental programs through the Hippo pathway were reported to function via phase separation ^32,33^. In addition, archetypical TFs with distinct sequence properties — including Myc, p53, and the well-studied Gcn4 from yeast — phase separate with Mediator complex or Mediator subunits ^3^. Clusters of Mediator, RNA polymerase II, and other components of the transcriptional machinery can form via phase separation and result in productive gene transcription ^5,6^. Similarly, several repressive regulators, such as HP1α, PRC2, MeCP2, and NELF also phase separate, and this state is thought to contribute to gene silencing ^16-20,34^. Accumulating evidence also suggests that Pol I–mediated transcription of ribosomal RNA is mediated by phase separation ^35^. Hence, multiple lines of evidence support a role for phase separation in regulating gene transcription.

Do these observations unequivocally demonstrate that phase separation is necessary for transcription? Some of the evidence is correlative, such as the effects observed using loss-of-function mutants. Efforts to probe the question using orthogonal approaches yielded mixed results. For example, prion-like low-complexity domains of fusion oncoproteins were overexpressed and formed transcriptionally repressive condensates ^25^. These observations were taken as evidence against the function of phase separation in transcription, but a key caveat is the dominant-negative effects of those domains, which limits insights into the function of transcriptional condensates. Another study varied TF expression levels and compared activity in cells with or without microscopically detectable condensates ^26^. Convincing differences were not observed. However, over-reliance on the LacO array confounded the assignment of the threshold level for the formation of condensates on chromatin. By contrast, titration of the muscle-specific TF MyoD showed a clear correlation between the threshold concentration for phase separation, the ability to activate transcription, and the phenotypic conversion of cells into cardiomyocytes ^36^, making a strong case for the role of phase separation for this specific TF in cell differentiation.

A key criticism of the phase separation model is the lack of rigorous comparison to previous quantitative models of transcriptional function ^21,22,37^. Importantly, the existence of multivalent interactions does not mean that they mediate function through phase separation. The interactions of two yeast TFs, Gcn4 and Gal4, with Mediator subunit Med15 are multivalent and dynamic ^38,39^, two key characteristics of phase-separating systems. However, these types of interactions also give rise to soluble higher-order complexes, presumably at concentrations below those in which they undergo phase separation with Mediator. The formation of soluble higher-order complexes below the saturation concentration, also called pre-percolation clusters, is typical for associative macromolecules that undergo phase separation ^28,40^, and they may therefore mediate transcription in the absence of phase separation. Head-to-head comparisons of the two models are therefore needed.

Mutations that change phase behavior are often used to correlate the driving force for phase separation with gene regulatory function ^3,23,30-32^. For instance, all aromatic residues in the transactivation domain of Gcn4 were replaced with alanine residues to attenuate phase separation ^3^. However, even less drastic mutations to Gcn4 can abrogate Med15 binding and the formation of potentially transcription-competent soluble complexes ^41^. Thus, the alanine substitutions may affect the formation of active species in both the “soluble complex” and the “phase separated condensate” models (**Fig. 1A**). Therefore, such blunt mutagenesis experiments cannot conclusively clarify the need for phase separation, or lack thereof, in transcriptional regulation. Recent work on N-Myc used a small molecule–based chemogenetic tool to induce phase separation and compare transcriptional activity in cells with and without phase separation at identical protein levels ^42^. The results showed that approximately 97% of N-Myc-regulated genes are not affected by phase separation, highlighting that transcription can be activated effectively by soluble complexes of TFs and transcriptional machinery. Nevertheless, the expression of 3% of genes was compromised without phase separation, arguing that the role of phase separation should be assessed.

**Figure 1.**
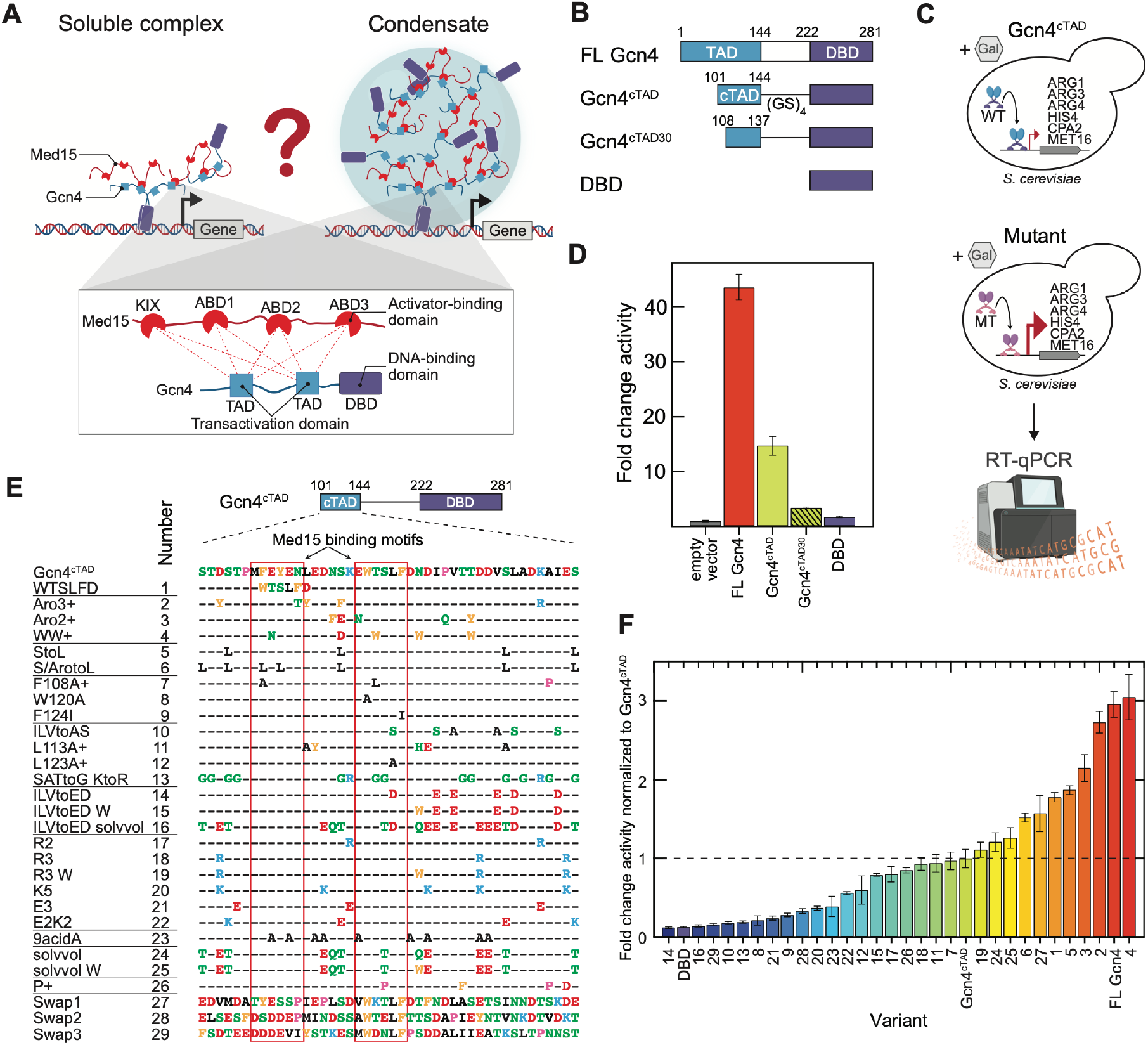
Gcn4 variants with a large range of activities can be generated. **(A)** Current models that may explain Gcn4 transcription factor activity, including soluble complexes (left) or condensates (right) formed by Gcn4, DNA, and Mediator. Both models are characterized by multivalent interactions between the Mediator and the transactivation domain of Gcn4. **(B)** Schematic of full-length (FL) Gcn4 and shorter variants, which contain transactivation domains and a DNA-binding domain (DBD). Med15 has a KIX domain and 3 activator-binding domains (ABDs). **(C)** Transcript levels of six genes activated by Gcn4 in a Gcn4 knockout strain with a galactose-responsive promoter were determined using RT-qPCR. **(D)** Activity mediated by the Gcn4 variants for six endogenous genes in (C), normalized to empty vector. At least three biological replicates per construct were measured with error bars indicating ± SEM. **(E)** Gcn4^cTAD^ variants designed for this study which: replace a Med15 binding motif with a stronger one (var 1); add aromatic (vars 2–4) or aliphatic residues (vars 5–6), respectively; replace aromatic (vars 7–9) or aliphatic residues (vars 14–16), respectively; added or removed charged residues (vars 17–22 and var 23, respectively); have increased side chain volume for some residues (vars 24–25); have added prolines (var 26); and have the same composition but different sequence patterning (vars 27–29). Many variants bear additional changes as indicated by + in the name; these TADs were used in previous work ^43^. **(F)** *In vivo* activities ordered from low to high activity for all Gcn4 variants relative to that of Gcn4^cTAD^. Horizontal line indicates WT Gcn4^cTAD^ activity. At least three biological replicates per construct were measured with error bars indicating ± SEM.

Herein, we performed a head-to-head, quantitative comparison of the roles phase separation and soluble complexes play in mediating Gcn4-driven gene transcription. We designed new Gcn4 variants, leveraging current knowledge of Gcn4–Med15 interactions ^38,39^, previously measured activity of Gcn4 mutants ^43,44^, and our conceptual understanding of sequence-encoded phase behavior ^45,46^. Our exhaustive analyses reveal that separating the transcriptional functions emerging from condensate or soluble complex formation is not straightforward, and the extant literature should be interpreted with sufficient caution. However, variants with the highest affinities do not function to their full potential and instead have activities that reflect their abilities to phase-separate with Med15. These data support the view that condensates indeed form in cells and determine activities. Furthermore, counter to the prevailing models, we show that multivalent DNA interactions interfere with Gcn4 phase separation because they promote the formation of soluble complexes. Our study reconciles opposing views in the literature and provides a transferable framework to critically evaluate the functional roles of other phase-separating systems.

## Results

### Gcn4 variants span a large range of activities

We generated a set of Gcn4 TAD variants to test whether the formation of soluble complexes or condensates account for its transcriptional activity (**Fig. 1A**). We introduced the variants into a Gcn4 construct, Gcn4^cTAD^, that has a previously described minimized TAD, with only 44 residues ^43^, and its native DNA-binding domain (DBD) (**Fig. 1B**). We generated a Gcn4 knockout yeast strain, expressed the Gcn4^cTAD^ construct from a plasmid under the control of the galactose-inducible *GAL1* promoter, and monitored transcripts from six endogenous target genes for Gcn4 by RT-qPCR (**Fig. 1C**). Those six genes have promoters with distinct arrangements of Gcn4 binding sites (**Fig. S1A**), which occur in different states of nucleosome occupancy and are bound by Gcn4 in response to physiological cues ^47^ (**Fig. S1B**). The wild-type (WT) Gcn4^cTAD^ led to ∼15-fold activation of the Gcn4 target genes compared to the empty vector (**Fig. 1D**). As controls, we expressed the full-length (FL) Gcn4, which yielded ∼43-fold activation; a further minimized Gcn4 construct with a 30-residue TAD, Gcn4^cTAD30 44^, which only yielded ∼3-fold activation; and the DBD alone, which behaved similarly to the empty vector. The six genes responded similarly, giving rise to a robust readout of gene activation.

Guided by the stickers-and-spacers framework ^45,46,48^, which treats phase-separating biomolecules as associative polymers with strongly and weakly adhesive elements (i.e., stickers and spacers), we rationally designed and generated 29 variants in this Gcn4^cTAD^ context with the following modifications (where + indicates that additional residues were mutated to mirror previously used variants ^43^; **Fig. 1E**):

Variants were designed to add a strong Med15 binding motif (var 1); to increase number of aromatic (vars 2–4) or aliphatic (vars 5–6) residues, as these are known to be determinants of transcriptional activity ^43,44,49,50^, mediate networking interactions in condensates formed by IDRs ^45,46^ and likely enable interactions with coactivators; to replace the native aromatic (vars 7–9) or aliphatic (vars 10–13) with alanine or charged residues, the latter of which would alter their solubilities ^46^. In addition, we added charged residues (vars 14–22) or replaced the native charged residues with alanine (var 23); replaced small solubilizing residues with larger ones (vars 24–25), or introduced proline residues (var 26) as helix breakers. Finally, we maintained sequence composition but changed its patterning (vars 27–29). We measured the transcripts of the target genes in response to expression of the variants, along with WT Gcn4^cTAD^, FL Gcn4 and the DBD as reference points.

We note that some of the mutations introduced are known to reduce Gcn4 binding to Med15, and indeed var 8 (W120A), var 9 (F124I) and var 12 (L123A+), all previously studied mutants, reduced transcriptional activation of the target genes. Variants with additional aromatic or aliphatic residues (vars 2–6, i.e., Aro3+, Aro2+, WW+, StoL, S/ArotoL) showed increased activation (**Fig. 1F**). Overall, 10 variants were more active than Gcn4^cTAD^ and 19 variants were less active, with activity values spanning a 30-fold range. Moreover, the transcriptional activities of previously reported Gcn4 variants correlated well with the activities measured in our studies (**Fig. S3**). These results validate our systematically designed set of Gcn4^cTAD^ variants with a wide range of sequence properties and transcriptional activities, which thus enables a more detailed assessment of the types of assemblies that mediate transcriptional activity *in vivo*.

### Homotypic phase separation of Gcn4^cTAD^ variants does not predict their ability to mediate transcriptional activity

To correlate the transcriptional activity of Gcn4 variants with defined properties *in vitro*, we purified all 29 Gcn4^cTAD^ variants (**Fig. S4A**), Gcn4^cTAD^, FL Gcn4, and DBD.

We first measured their propensity to undergo homotypic phase separation. DBD alone did not phase separate under any experimentally accessible conditions (**Fig. S4B**), whereas Gcn4^cTAD^ and FL Gcn4 formed condensates (**Fig. 2A**) above threshold concentrations, also known as saturation concentrations (*c*_sat_), in the presence of molecular crowders. We determined their saturation concentrations by separating the dilute and dense phases and measuring the dilute phase concentrations using analytical HPLC (**Fig. 2B**). We found that the saturation concentration of Gcn4^cTAD^ was ca. 2.5 times higher than that of FL Gcn4 (**Fig. 2C**). These data agree with previous results that the TAD mediates phase separation ^3^ and that multivalence is often encoded in a uniformly distributed manner across the sequence ^46,51^, resulting in a higher drive for phase separation for FL Gcn4.

**Figure 2:**
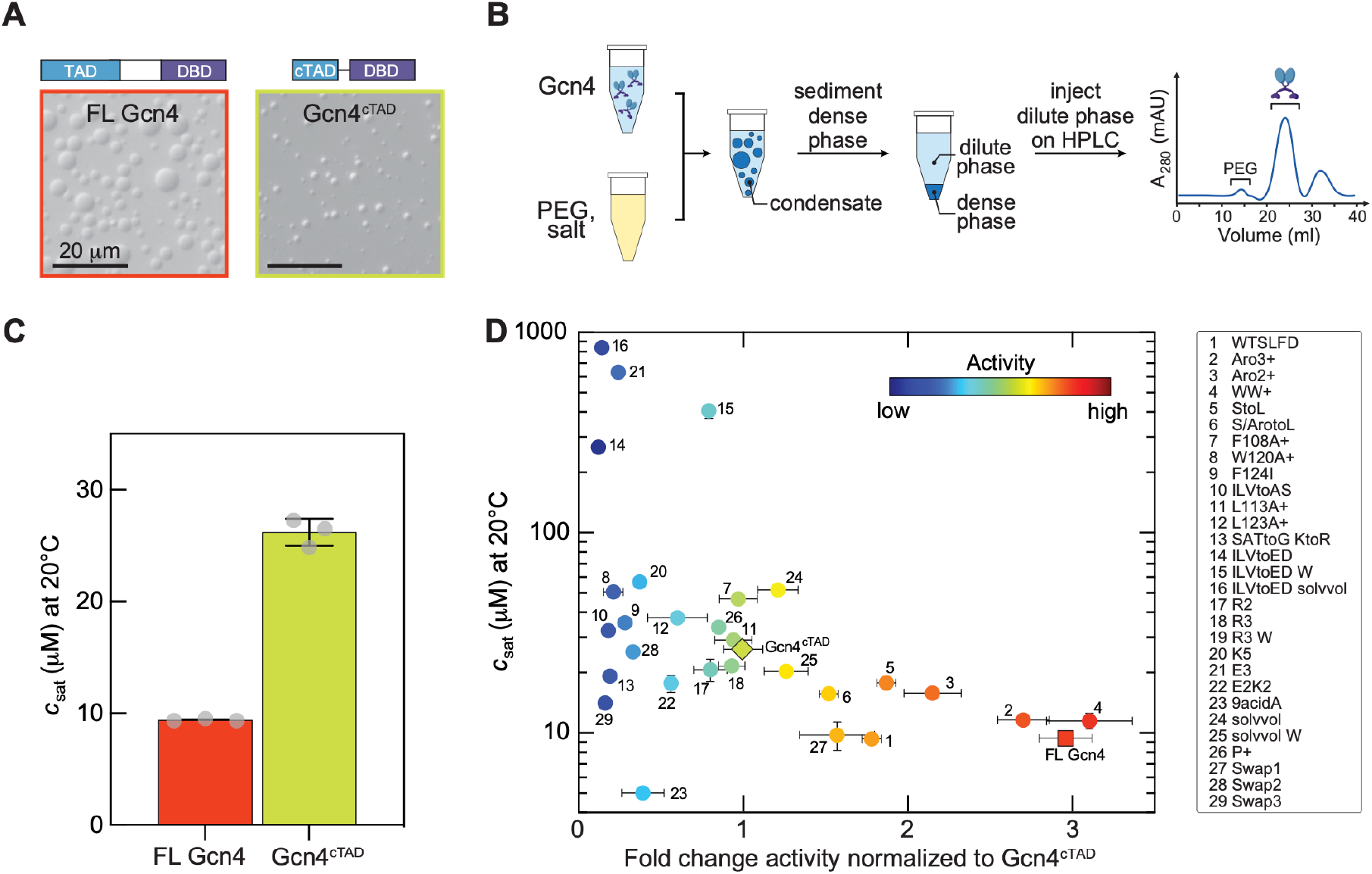
Homotypic phase separation propensity does not predict Gcn4-mediated transcriptional activity. **(A)** DIC images of FL Gcn4 and Gcn4^cTAD^ showing that both proteins form condensates in the presence of 10% PEG 8K *in vitro*. **(B)** Schematic showing the approach with which the saturation concentration (*c*_sat_) of Gcn4 and its variants was determined using analytical HPLC^52^. Phase separation was induced by adding salt and PEG to the protein solution, and the solution was centrifuged to separate the dense and dilute phases. The protein and PEG concentrations in the dilute phase were determined using analytical HPLC (see Methods). **(C)** The saturation concentrations (*c*_sat_) for FL Gcn4 and Gcn4^cTAD^. Individual data points from n = 3 measurements are indicated, with error bars indicating ± SEM. **(D)** Saturation concentrations of all Gcn4 variants as a function of their transcriptional activities relative to that of Gcn4^cTAD^. We were not able to measure a saturation concentration for var 19. Error bars indicate ± SEM from at least three independent measurements. Solution conditions were 20 mM HEPES (pH 7.3), 150 mM potassium acetate, 10% PEG 8K, 2 mM DTT.

We then evaluated the driving force for phase separation for all 29 Gcn4^cTAD^ variants and compared that to the observed transcriptional activities. All variants formed condensates in the presence of the molecular crowder PEG 8K (**Fig. S4B**), with saturation concentrations that varied by 2.5 orders of magnitude (**Fig. 2D**). Variants with additional aromatic or aliphatic residues (vars 1–6) displayed lower saturation concentrations relative to WT Gcn4^cTAD^, mirroring their higher transcriptional activities. Conversely, reducing the number of aromatic residues (vars 7–9) or replacing aliphatic residues with charged ones (vars 14–16) led to weakened phase separation and lower transcriptional activities. While these results are suggestive of a trend whereby the driving force for phase separation roughly reflects the respective activities, several variants that mediated reduced activities did not show markedly altered saturation concentrations relative to WT Gcn4^cTAD^ (vars 10, 13, and 28). Moreover, some transcriptionally inactive variants displayed low saturation concentrations (e.g., vars 23 and 29) that were similar to those of the most active variants. In sum, our compendium of Gcn4 variants demonstrates that the propensity for homotypic phase separation does not reliably predict transcriptional activities *in vivo*.

### DNA binding promotes dissolution of Gcn4^cTAD^ condensates

Recent investigations propose models wherein multiple DNA sites serve as scaffolds that facilitate the formation of TF condensates ^2,10-12^. Based on these prevailing models, we hypothesized that binding to multiple cognate DNA sites would enhance the driving force for phase separation for Gcn4^cTAD^. To test this hypothesis, we generated DNA constructs of identical length and containing one, two, or four Gcn4 response elements (GREs), each of which accommodate a homodimer of Gcn4 (**Fig. 3A**). Unexpectedly, titration of the DNA constructs into phase-separated Gcn4^cTAD^ samples progressively weakened phase separation, resulting in complete dissolution of the Gcn4^cTAD^ condensates at stoichiometric levels (**Fig. 3B**). Notably, the DNA constructs with a higher number of GREs dissolved condensates at lower concentrations, unambiguously demonstrating that Gcn4^cTAD^ phase separation is weakened by multivalent DNA binding.

**Figure 3:**
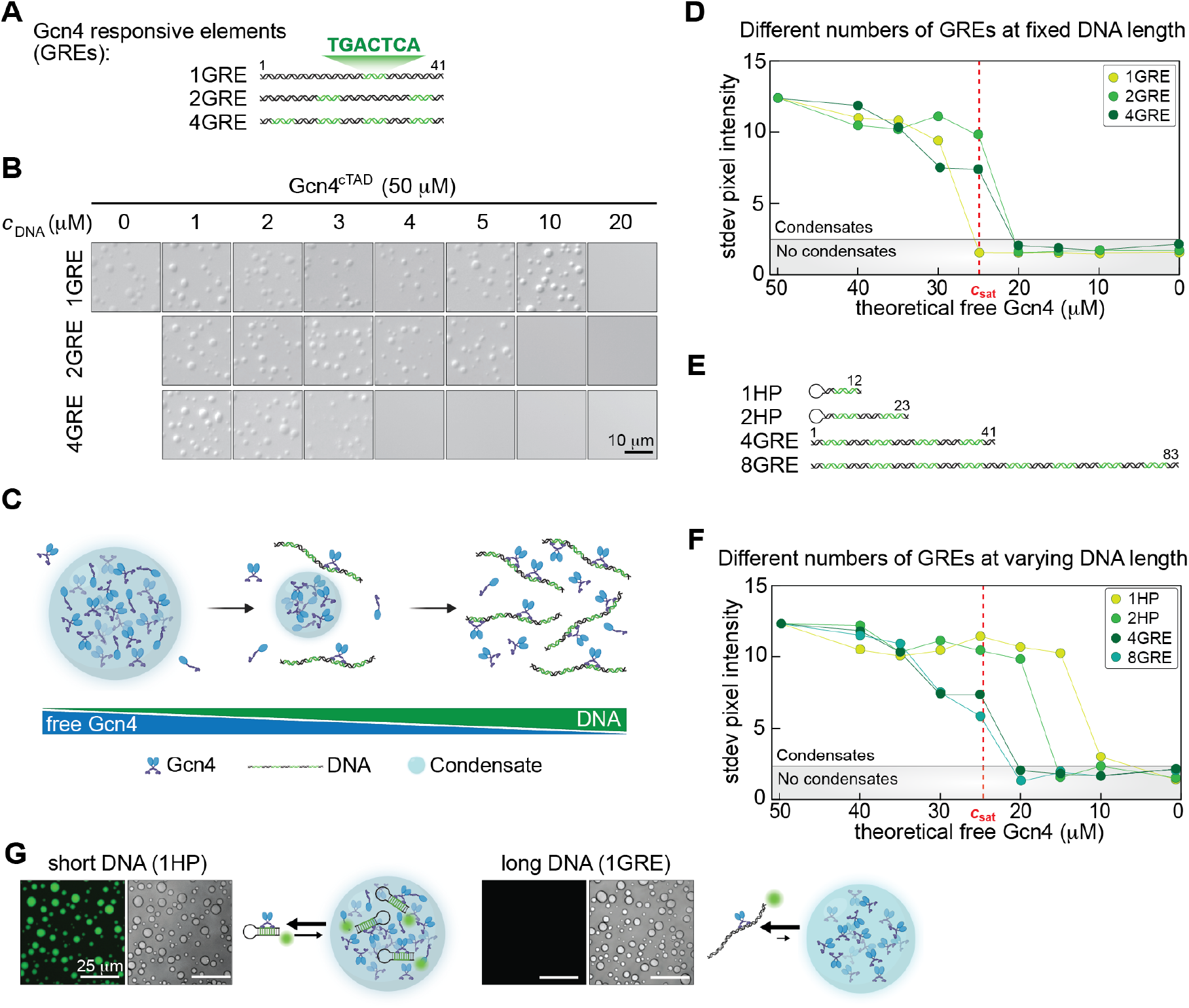
DNA with multiple binding sites for Gcn4 suppresses phase separation. **(A)** Schematic of DNA constructs with fixed length and varying numbers of Gcn4 responsive elements (GREs). **(B)** DIC images of Gcn4 solution (50 μM) with increasing concentrations of DNA. **(C)** Schematic of a model in which free Gcn4 phase separates above a saturation concentration and binding to DNA solubilizes Gcn4. **(D)** Index of dispersion *vs* the theoretical free Gcn4 concentration after stoichiometric binding to DNA [*c*_Gcn4_ – (2 × #GRE × *c*_DNA_)] for DNA constructs of the same length but varying GREs. The dashed red vertical line indicates the saturation concentration of Gcn4^cTAD^. **(E)** Schematic of DNA constructs with varying numbers of GREs and length. Short constructs are formed by single-stranded DNA with hairpins for stability reasons. **(F)** Index of dispersion *vs* the theoretical free Gcn4 concentration after stoichiometric binding to DNA with varying number of GREs and length. **(G)** DIC and fluorescent micrographs of solutions of Gcn4^cTAD^ and short or long DNA. 1HP and 1GRE DNA constructs were labeled with Alexa-Fluor-488 at the 5’-end.

Our observations indicate that Gcn4^cTAD^–DNA complexes are more soluble than unbound Gcn4^cTAD^. The data support a model in which binding of Gcn4^cTAD^ dimers to DNA (i.e., two protein molecules per GRE) reduces the amount of unbound protein available for phase separation (**Fig. 3C**). According to this model, the concentration of free Gcn4^cTAD^ must drop below the saturation concentration for condensate dissolution. To test this model, we collected microscopy images across a broader range of DNA concentrations and plotted the standard deviations of the pixel intensity histograms (which fall below a baseline level in the absence of microscopically detectable condensates) against the expected concentrations of unbound Gcn4^cTAD^ molecules. We assumed stoichiometric binding because the binding affinity of the DBD for DNA is in the nanomolar range, which leads to high fractional occupancy in concentration regimes used for condensate formation (see **Fig. S5**). The data for all DNA constructs collapsed within error onto a single curve, supporting the model that Gcn4^cTAD^–DNA complexes are soluble and therefore less disposed to undergo phase separation (**Fig. 3D**).

Next, we hypothesized that the source of the solubility of the protein–DNA complexes was their high net charge. To test this, we generated a series of DNA constructs with varying numbers of GREs and constant linker length, so the resulting DNA constructs varied considerably in length (from 12 to 83 bp) (**Fig. 3E**). For these constructs, the simple model did not adequately describe the data (**Fig. 3F, Fig. S6**), i.e., the shorter DNA molecules were not as efficient at solubilizing condensates as the longer ones, pointing to the possibility that Gcn4^cTAD^ complexes with short DNA constructs were weakly permissive to phase separation. Indeed, short DNA constructs partitioned into the condensates more effectively than long DNA constructs (**Fig. 3G**). By contrast, long DNA molecules, which enhance Gcn4^cTAD^ solubility rather than its phase separation, do not partition into homotypic Gcn4^cTAD^ condensates under our experimental conditions.

These observations challenge the widely held assumption that multiple DNA binding sites should serve as a scaffold for phase separation. Instead, our results demonstrate that multivalent interactions alone are not sufficient to mediate phase separation. If the resulting higher-order complexes are highly soluble, multivalent interactions may well counteract innate propensities for phase separation.

### The Mediator subunit Med15 rescues phase separation of soluble Gcn4–DNA complexes

If transcription is mediated by biomolecular condensates, the transcriptional machinery recruited by the TF may aid in condensate formation. Indeed, TFs are known to co-phase separate with the Mediator complex ^3,4^. In particular, Gcn4 co-phase separates with Med15, a key subunit in the Mediator complex that mediates Gcn4 function ^3^. Med15 contains three activator-binding domains (ABD) and one KIX domain that interact with the cTAD of Gcn4 ^39^ (**Fig. 4A**).

**Figure 4:**
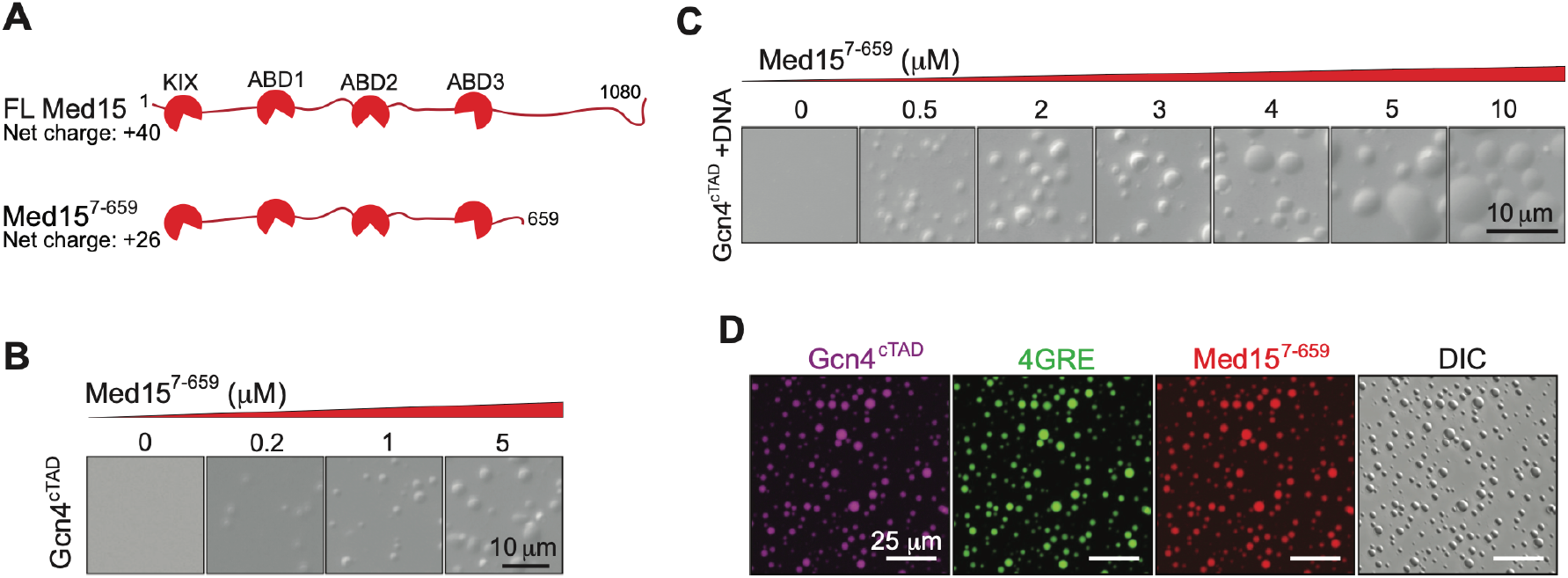
The Mediator subunit Med15 rescues Gcn4/DNA phase separation. **(A)** Schematic of Med15, which is part of the tail module of the Mediator complex^53^. Med15 has three activator-binding domains (ABD) and a KIX domain that have been shown to interact with Gcn4^cTAD 54^. **(B)** DIC images of 50 μM Gcn4^cTAD^ solutions with increasing concentrations of Med15^7-659^. **(C)** DIC images of solutions containing 50 μM Gcn4^cTAD^ and 2.5 μM 4GRE DNA with increasing concentrations of Med15^7-659^. **(D)** Co-localization of Gcn4, Med15, and 4GRE DNA oligonucleotide in condensates. Solution conditions were 20 mM HEPES (pH 7.3), 150 mM potassium acetate, 2 mM DTT.

We expressed and purified a previously used, truncated version of Med15 (Med15^7-659^) ^38^ (**Fig. S4**), which includes all four interaction domains, but lacks the C-terminal domain, which is required for incorporation into the Mediator complex. Titrating the truncated Med15 into solutions of Gcn4^cTAD^ enhanced Gcn4^cTAD^ phase separation (**Fig. 4B**). The same was observed for FL Gcn4 (**Fig. S7A**). We also observed phase separation of Med15 alone at concentrations above 5 μM in the presence of crowders (**Fig. S7B**).

While Med15 was not previously reported to bind DNA, we observed that Med15 formed condensates with DNA (**Fig. S7C**). Notably, Med15 was able to rescue the phase separation propensity of soluble Gcn4^cTAD^–DNA^4GRE^ complexes (**Fig. 4C**). Co-condensation of Gcn4^cTAD^ and Med15 showed only a weak dependence on DNA concentration, i.e., condensates were neither dissolved nor stabilized by the presence of DNA (**Fig. S7D**). We used fluorescence microscopy to confirm that Gcn4^cTAD^, Med15, and DNA^4GRE^ co-localize in condensates (**Fig. 4D**). Taken together, our observations support a model wherein the positive net charge of Med15 offsets the high negative net charge of Gcn4^cTAD^–DNA complexes and facilitates phase separation over large Gcn4/DNA concentration ranges. Thus, binding to Mediator may facilitate the formation of Gcn4 condensates on regulatory DNA sites in the genome.

### Quantification of Med15 incorporation into Gcn4^cTAD^ variant condensates

Next, to test the phase separation model, we investigated the extent to which different Gcn4 variants recruit Med15 into condensates. Their co-phase separation propensity provides a readout to compare the *in vitro* phase separation properties of each variant with the corresponding transcriptional activity they mediate *in vivo*. Our Gcn4 variants include those harboring mutations in Med15 binding motifs, having altered net charge or charge distributions, or bearing additional hydrophobic residues that strengthen engagement with Med15 (see **Fig. 1E**). These perturbations should provide a clear rationale for why their phase behavior with Med15 would differ from their homotypic phase behavior.

Using fluorescence microscopy, we quantified the level of Med15 incorporation into 30 Gcn4^cTAD^/DNA^4GRE^ condensates those formed using the DBD and full-length Gcn4 (**Fig. 5A**). 10% of Med15 in each sample was fluorescently labeled to enable its quantification. We chose constant concentrations for Gcn4^cTAD^ variants, Med15, and 4GRE DNA, to enable quantitative comparisons of the phase behavior of the variants. We found that different variants formed different-sized condensates with DNA and Med15, and that Med15 partitioned into these with varying efficiencies (**Fig. 5B**). Var 23 (with acidic residues replaced with alanine) was the only variant that showed obvious stronger partitioning into relatively smaller condensates (**Fig. 5B**). These observations reflected the differing abilities of Gcn4 variants to incorporate Med15. We therefore quantified the fraction of Med15 incorporated into condensates of each variant by determining the partition coefficients (i.e., the relative concentrations of Med15 in the dense vs. the dilute phases (**Fig. 5C**)) and the dense phase volume fractions (**Fig. 5D**). The incorporated fraction of Med15 varies by about two orders of magnitude for condensates formed by different Gcn4 variants (**Fig. 5E**).

**Figure 5:**
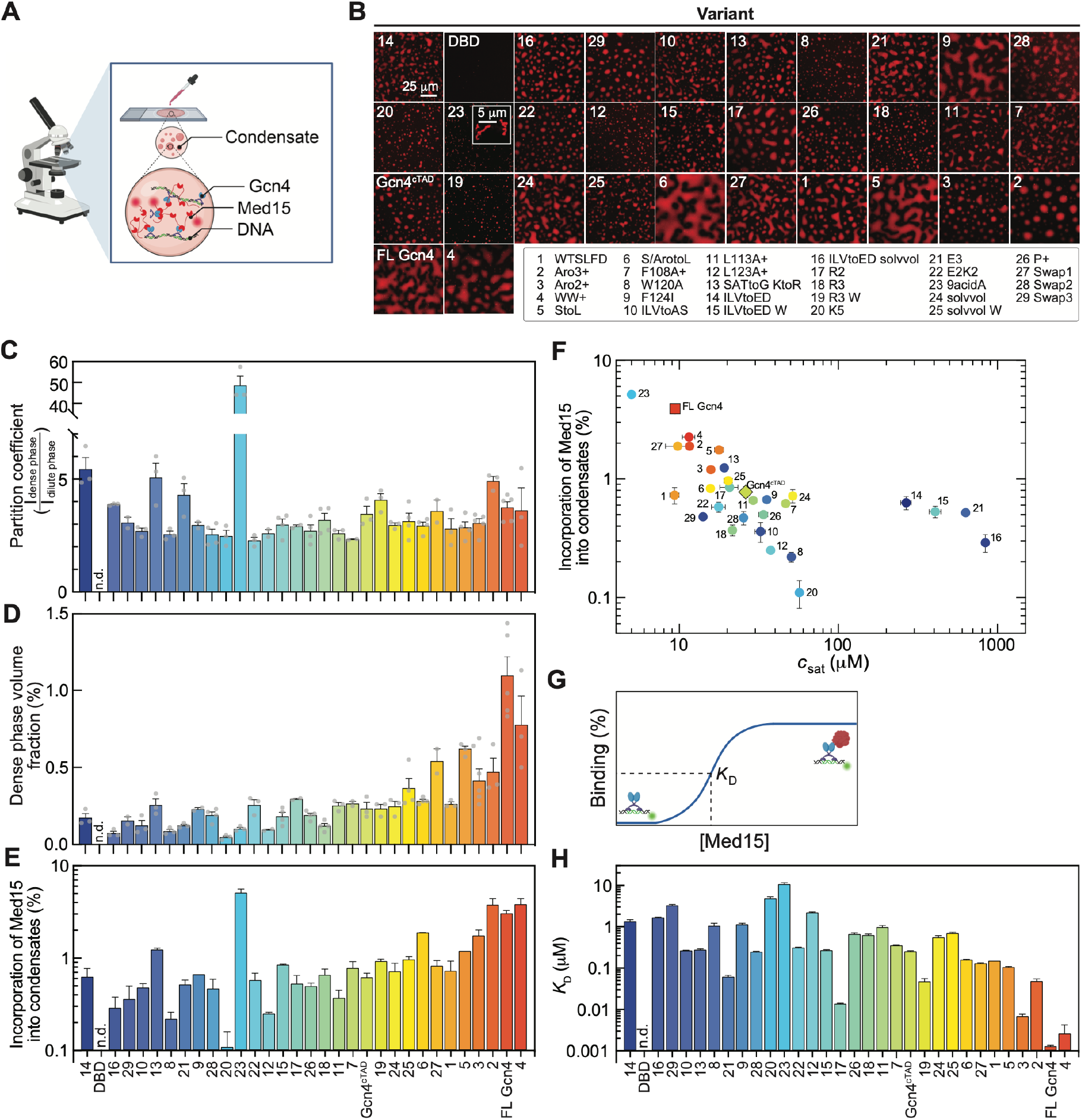
Does the formation of soluble complexes or condensates explain the observed Gcn4 transcriptional activity? **(A)** Schematic showing condensates formed by 50 μM Gcn4^cTAD^ variants, 10 μM Med15 (10% N-terminally labeled with LD-555), and 0.5 μM 4GRE DNA constructs. The fraction of Med15 incorporated into condensates for all Gcn4 variants was determined by taking z-stack images via confocal microscopy. **(B)** Confocal microscopy images showing condensates formed by Med15, 4GRE DNA, and all Gcn4 variants, with variants ordered by activity. **(C)** Partition coefficient of Med15^7-659^ into Gcn4^cTAD^/4GRE condensates and **(D)** dense phase volume fraction of these condensates. **(E)** Incorporation of the fraction of Med15 into condensates across all Gcn4 variants, with variants ordered from low to high activity. Error bars indicate ± SEM from at least three independent measurements. We did not observe any condensates when adding DBD to Gcn4^cTAD^/4GRE (indicated by n.d.). **(F)** Med15 recruitment into condensates as measure of heterotypic phase separation vs homotypic saturation concentrations of Gcn4 variants. **(G)** Schematic for determination of the affinity of Med15 to the Gcn4-DNA complex using fluorescence anisotropy binding assays. **(H)** Dissociation constant *K*_D_ plotted against all Gcn4 variants with variants ordered by their activity values. The colors of the bars correlate with the activities as in Fig. 1F. Error bars indicate ± SEM from at least three measurements. Solution conditions were 20 mM HEPES (pH 7.3), 150 mM potassium acetate, 2 mM DTT.

We compared the homotypic saturation concentrations and the extent of incorporation of Med15 into condensates (heterotypic phase separation) of the Gcn4 variants with DNA and Med15 (**Fig. 5F**). While the two properties were strongly correlated for many variants, they show different patterns for others. Vars 14, 15, 16, and 21 have a more negative net charge per residue than other variants, which weakens their homotypic phase separation but seems to stimulate co-phase separation with DNA and Med15. Variants 1, 6, 7, 9, 17, and 24 show similar abilities to incorporate Med15 as Gcn4^cTAD^ but vary strongly in their homotypic saturation concentrations. The behavior of these variants contributed to the relatively poor predictive power of the homotypic saturation concentrations towards the *in vivo* activities. Therefore, the different abilities of Gcn4 variants to incorporate Med15 may provide an explanation for their transcriptional activities.

### Med15 forms soluble complexes of various strengths with DNA-bound Gcn4^cTAD^ variants

The multivalent interactions between Gcn4, DNA, and Med15 mediate phase separation, but they also mediate the formation of soluble complexes ^38^. Such soluble complexes or pre-percolation clusters exist below the saturation concentration and coexist with condensates ^28,55^. To test whether these soluble complexes may be the molecular species that activate transcription, we performed fluorescence anisotropy binding assays to determine the affinities of Med15 to Gcn4– DNA complexes (**Fig. 5G, Fig. 5C,D**).

The resulting solutions only contained soluble complexes, not condensates (**Fig. S5C**) as expected from the low concentrations of Gcn4 used and as confirmed by microscopy. The affinities of Med15 for the Gcn4 variant–DNA complexes varied by four orders of magnitude (**Fig. 5H**). While a general relationship between activity and affinity seemed evident, there was significant variation in affinities between variants with low activities (e.g., vars 8, 21, and 26).

### Med15 incorporation into condensates and formation of soluble complexes predicts Gcn4 variant–mediated activity

Which of the two transcriptional models can best explain the *in vivo* transcriptional activities of the Gcn4 variants? To answer this question, we assessed how the two biophysical readouts tracked with the activities of different Gcn4 variants. We plotted Med15 incorporation into condensates against transcriptional activities and found a predictive trend, where stronger incorporation tracks with higher activity (**Fig. 6A**). Similarly, the affinities (*K*_D_ values) of the Gcn4–DNA complexes for Med15 increase with the activities of variants in a predictive manner (**Fig. 6B**). The direct comparison of the two biophysical readouts shows that they are highly correlated (**Fig. 6C**). These observations point to the fact that the interactions mediating the formation of soluble complexes and condensates strongly overlap.

**Figure 6:**
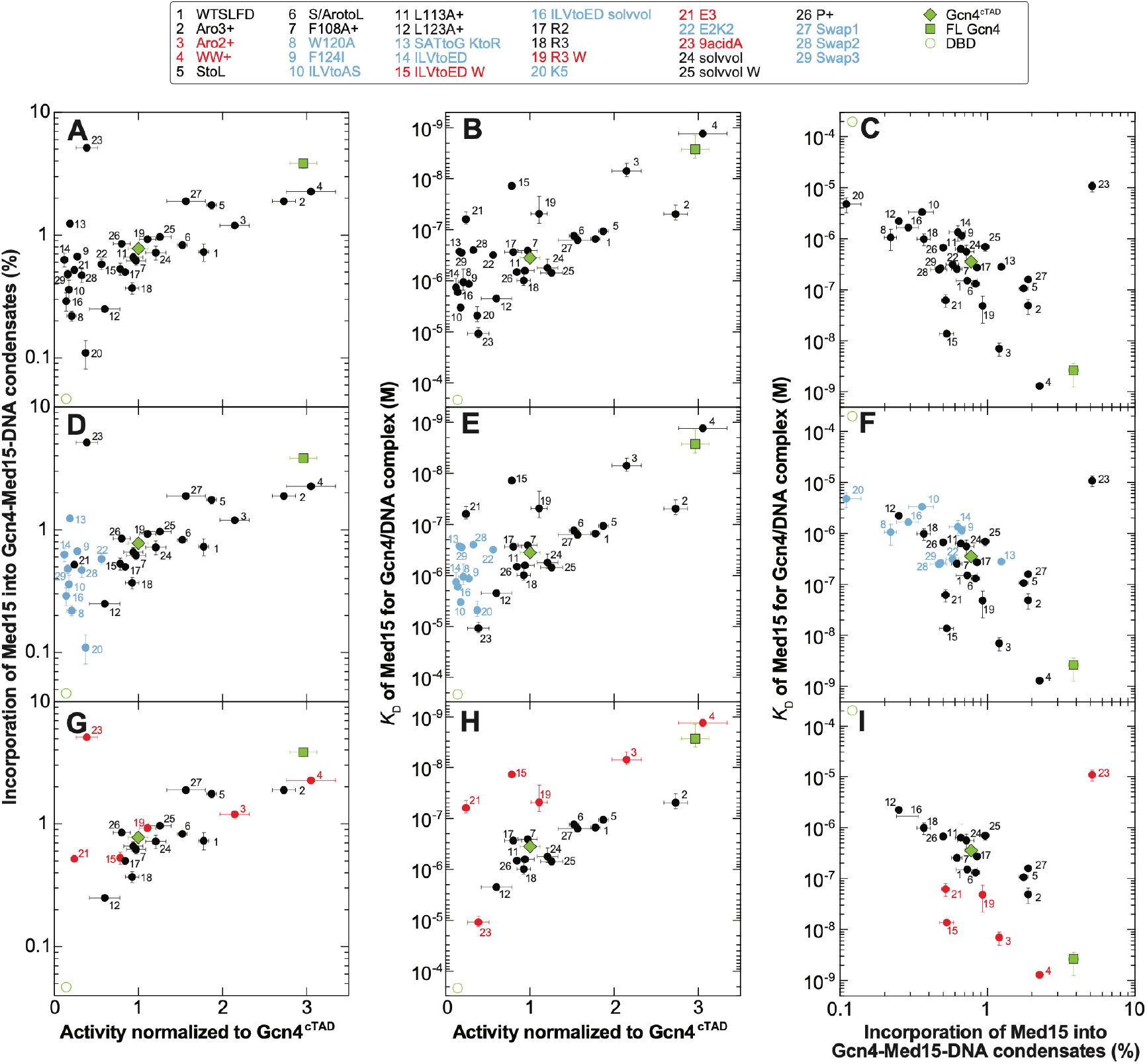
Incorporation of Med15 into Gcn4-DNA-Med15 condensates and binding to Gcn4/DNA predict transcriptional activity. Comparison of how well the phase separation model **(A, D, G)** and soluble complex model **(B, E, H)** explain Gcn4 activity. **(C, F, I)** Comparison of the two biophysical readouts. Solution conditions were 20 mM HEPES (pH 7.3), 150 mM potassium acetate, 2 mM DTT. WT Gcn4^cTAD^ is represented by a diamond, FL Gcn4 by a square, and DBD by an open circle. The DBD is shown as an open circle with approximate position because no binding of Med15 to preformed DBD-DNA complex was detected up to 300 mM of Med15, and we observed no condensates for DBD/DNA/Med15 mixtures.

The set of three plots revealed a set of variants with low transcriptional activities (shown in blue in **Fig. 6D–F**) but a large range in their abilities to incorporate Med15 into condensates (**Fig. 6D**) and to interact with Med15 in soluble complexes (**Fig. 6E**). Thus, the predictive relationship between activities and biophysical properties did not hold for these variants, although the biophysical properties themselves remained highly correlated (**Fig. 6F**).

We considered whether the mutations in these variants could perturb not only their abilities to interact or phase separate with the transcriptional machinery but also other biological behaviors such as their abundance or subcellular localization. If certain Gcn4 variants were mislocalized, they would not elicit transcriptional activities *in vivo*, even if their abilities to interact with Med15 *in vitro* would suggest otherwise. Therefore, we used confocal fluorescence microscopy to examine the subcellular distributions of a curated set of variants, which were expressed as GFP fusions (**Fig. S9A–B**). Cells expressing WT Gcn4^cTAD^ or the DBD had stronger GFP signals in the nucleus than in the cytoplasm, as expected (**Fig. S9B**). Quantification of the total fluorescence intensity in the nucleus over that in the cytoplasm showed values of ∼0.7–0.8 (**Fig. S9C**). Given the clear nuclear accumulation of these variants, these values reflect the smaller relative area occupied by the nucleus in 2D confocal images. Var 8, a variant with low activity and low ability to interact with Med15, showed the strongest enrichment in the nucleus, implying that its low activity was likely not driven by mislocalization. For var 29 and var 13, the ratios of nuclear vs cytoplasmic protein levels were lower than for Gcn4^cTAD^ (**Fig. S9C**), indicating that the mislocalization of these variants might contribute to their low activity.

Next, we examined the enrichment of the same set of Gcn4 variants at gene promoters by performing ChIP experiments. WT Gcn4^cTAD^, the DBD-only construct, the low-activity–promoting var 8, and the strongly activating variant 3 show similar enrichments at promoters of Gcn4-responsive genes. By contrast, other variants, such as var 4, mediate high activity despite low enrichment at promoters (**Fig. S9D**). The relatively lower enrichment of variants such as var 15, relative to that of WT Gcn4^cTAD^, thus does not explain their unexpectedly low activities. We note that ChIP relies on unimpeded access of the antibody to the epitope. It is also possible that other biological factors that are difficult to identify or quantify may also be at play.

The correlation between the two biophysical readouts for this group of poorly active variants was very strong and on par with those of highly active variants (**Fig. 6F**). We thus concluded that the low transcriptional activity of this group of variants is due to biological factors that are not captured by our biophysical measurements.

### Partial separation-of-function variants reveal the role of phase separation

For a subset of variants (shown in red in **Fig. 6G–I**), the correlation between the two biophysical readouts was poorer than for the other variants. These variants had additional aromatic residues (vars 3, 4, 19, and 21) or aspartate residues (vars 15 and 21), which are features known to encode strong transcriptional activity in TADs ^43,49^. The abilities of these variants to incorporate Med15 into condensates was on par with what would be expected based on their *in vivo* activities (**Fig. 6G**). However, their affinities for Med15 were higher (lower *K*_D_ values) than would be expected based on their activities (**Fig. 6H**). Thus, this set of variants (shown in red) may accomplish partial separation of function between the interactions that drive the formation of soluble complexes vs condensates. Var 23, in which all acidic residues are mutated to alanine residues, was the only notable outlier — this mutant is biologically inactive but differs from the inactive variants shown in blue because of its distinct effects on phase separation and binding. Var 23 incorporates Med15 very strongly into condensates (as also visible in **Fig. 5B, C, E**) but interacts with Med15 with weak affinity.

Do these variants suggest that soluble complexes or condensates explain activities better? Their phase separation propensities have a clear predictive relationship with in vivo activities, whereas their affinities for soluble Med15 is higher than expected (**Fig. 6K**). Our data thus suggest that these high-affinity variants form condensates in cells that determine their activities, but those are attenuated relative to the expectation from their affinities for Med15.

Together, our results show that the driving forces for the formation of soluble complexes and condensates are highly correlated. However, we also find that the formation of soluble complexes and condensates can be partially decoupled. The variants that break this strong coupling, i.e., those displaying affinities for Med15 that do not result in expected *in vivo* activities, reveal that the ability to phase separate with Med15 is intrinsically encoded in the sequence of Gcn4 and attenuates transcriptional activity *in vivo*.

## Discussion

Whether transcription is mediated via biomolecular condensates or non-stoichiometric soluble assemblies distinct from phase-separated condensates has been intensely debated in the literature ^8,21,22^, yet studies that compare the two models head-to-head via mutational interrogation have been missing. Here, we present strong evidence that the transcriptional activities of many Gcn4 variants are explained both by their abilities to phase separate with Med15 as well as by their abilities to form soluble complexes with the key coactivator Med15 of the Mediator complex. Hence, biophysical readouts from relatively simple *in vitro* systems can be predictive of *in vivo* activities if they reconstitute the major relevant interactions. Among the many unexpected findings with implications for extant data in the literature, we report that homotypic phase separation of Gcn4 variants does not explain their *in vivo* activities well. Furthermore, data from few mutants are likely insufficient for supporting one model over the other. Our results thus caution against the use of few sequence variants to correlate their driving forces for phase separation, whether homotypic or heterotypic, with their gene regulatory activities.

Another important finding cautions against the widely held assumption that binding to DNA serves to scaffold the formation of TF condensates. We demonstrate that, contrary to expectations, DNA binding increases the solubility of the TF Gcn4 and suppresses condensate formation. Additionally, multivalent engagement on DNA is insufficient for phase separation. Our results point to the importance of two factors for phase separation: the networking ability of the constituent molecules and the relatively low solubility of the resulting complexes. These results build on previous computational, experimental, and theoretical work ^46,56,57^ to highlight the underlying principle that condensates are formed by phase separation that is coupled to percolation (PSCP) ^27,58,59.^

Importantly, our careful, quantitative measurements show that interactions which drive the formation of condensates or soluble complexes are largely overlapping, and only extensive mutagenesis enabled us to partially deconvolute them. Indeed, the long-studied Gcn4 mutants and the plurality of our rationally designed variants display indistinguishable propensities to phase separate and form soluble complexes with Med15. The intertwined abilities to mediate phase separation and associate with coactivators on DNA thus seems to be programmed into the TAD sequence. Some of the known sequence features that are important for the activity of TFs include hydrophobic, particularly aromatic, and negatively charged residues ^43,49^. These sequence features can be reinterpreted in light of our results. The negative charge prevents phase separation of Gcn4 with DNA alone but enables phase separation with the positively charged Med15. The aromatic and aliphatic hydrophobic residues mediate interactions with Med15 and help mediate the network formation required for condensate formation. Similar relationships are expected for the interactions between TFs and other cognate coactivators.

Our ability to generate partial separation-of-function variants is anchored in the differences between the physical properties of soluble complexes and phase separated condensates, i.e., in the fact that condensates have emergent properties that small complexes do not possess ^27,59^. The variants in question, which displayed high affinities for Med15 in soluble complexes, failed to activate expression to the levels that would be expected. By contrast, their propensities to recruit Med15 into condensates better explained their transcriptional activities *in vivo*. When viewed collectively, our data reveals that both soluble complexes *and* condensates can drive transcription. However, high-affinity binding is not always beneficial, potentially because the high affinities give rise to condensates with strong internal percolation and attenuated transport processes, as has been previously demonstrated ^60^.

Overall, transcription happens in an environment rich in several types of condensates, including chromatin condensates ^16,17,61^, condensates of epigenetic regulators ^62^, and splicing condensates ^15^. Real time visualization of transcriptional condensates containing Mediator and RNA Polymerase II has recently shown that they approach promoters, transiently engage, and activate genes in sustained bursts ^63^. The ability of TFs to dynamically interact with such condensates may not require phase separation of the TFs themselves.

It is important to note that rather than ascribe to one side of the debate or the other, our results fit into an overall picture painted by recent work that highlights the role of condensates while also suggesting (1) little additional transcriptional activity is gained when TFs transition into a phase-separated state ^26,42^; and (2) little loss of activity is experienced when condensates are actively dissolved ^64^. However, specific genes experience direct regulatory effects due to phase separation, suggesting that the emergent properties of condensates can enable additional layers of target gene regulation and impart new biological functions. These may include changes to chromatin structure ^30,65,66^, partitioning of activating factors or exclusion of repressive factors ^67^, or changes to the electric potentials in the cell ^68,69^. The exact mechanisms by which transcriptional condensates exert their functions offer exciting new directions for future studies.

## Limitations of the study

Gcn4 variants were expressed from an exogeneous promoter at slightly super-physiological levels. Future work should compare the driving forces for phase separation with transcriptional activities of variants expressed at endogenous levels. We used the single Med15 subunit of the Mediator to reconstitute multicomponent condensates *in vitro* and omitted its disordered glutamine-rich tail that limits its solubility ^70^ and therefore creates technical limitations. Complex formation and the phase behavior of this reductionist system reflects cellular activity surprisingly well despite the effects other protein components can have on phase behavior. We interpret this as indication that Gcn4 and Med15 are the major nodes that determine transcription of Gcn4-dependent genes ^71^. Nevertheless, it will be interesting to reconstitute more complex transcriptional condensates *in vitro* and test whether their activity recapitulates transcriptional activity in yeast.

## Acknowledgements

We thank Melissa Marzahn, Ambuja Navalkar, and Joseph Brett for fruitful discussions. Research reported in this publication was supported by the National Institutes of Health under award number GM154414 (to T.M. and A.Z.A.) and NS108376 (to A.Z.A), the American Lebanese Syrian Associated Charities (ALSAC, to T.M. and A.Z.A.), the National Science Foundation award CEE:EFRI 02127 (to A.Z.A), and the St. Jude Research Collaborative on the Biology and Biophysics of RNP granules (to T.M.).

