## Supplementary Information for "Reconciling competing models on the roles of condensates and soluble complexes in transcription factor function"

Supplementary Methods
Supplementary Tables S1 – S4
Supplementary Figures S1 – S9
Supplementary References

### Supplementary Methods

*Design of Gcn4 activation domain variants and Med15 constructs*

The general sequence design of Gcn4 variants used in this study was based on Reference [1] and spans the central activation domain (residues 101-141) from *S. cerevisiae* (UniProt: P03069). The central activation domain is connected by a short $(GS)_4$-linker to the DNA-binding domain of Gcn4 (residues 222-281). We also included the full-length Gcn4 protein in our study (residues 1-281). For downstream fluorescence labeling purposes, serine 262 in the DNA-binding domain was substituted by a cysteine for $Gcn4^{cTAD}$ and FL Gcn4 (underlined in the sequences below). For Med15, a truncated sequence (residues 7-659) was used which includes all 4 activator-binding domains from *S. cerevisiae* (UniProt: P19659). This construct has previously been used in *in vitro* studies of Gcn4-Med15 interactions [2]. Additionally, a shorter construct of Gcn4 spanning just 30 residues was generated and referred to as $Gcn4^{cTAD30}$.

The coding sequences for the variants were synthesized (Genscript) including a coding sequence for an N-terminal TEV protease cleavage site and 5′ and 3′ attL sites for insertion into a ENTR vector and Gateway cloning. The sequences were recombined via LR reactions into the pDEST17 vector (ThermoFisher), which includes an N-terminal 6xHis-tag coding sequence. For in vitro experiments, the N-terminal 6xHis-tag was cleaved using the TEV protease cleavage site, leaving only an additional GS sequence at the N-terminus of each of the constructs (underlined in Table S1). Amino acid sequences of all constructs are shown in Table S1.

**Table S1: Sequences of $Gcn4^{cTAD}$ variants used in this study.**

| Name | Sequence |
|---|---|
| $Gcn4^{cTAD}$ (44-mer) | (GS) STDSTPMFEYENLEDNSKEWTSLFDNDIPVTTDDVSLADKAIESGSGSGSGSPESSDPAALK<br>RARNTEAARRSRARKLQRMKQLEDKVEELLCKNYHLENEVARLKKLVGER |
| F124I | (GS) STDSTPMFEYENLEDNSKEWTSLIDNDIPVTTDDVSLADKAIESGSGSGSGSPESSDPAALK<br>RARNTEAARRSRARKLQRMKQLEDKVEELLCKNYHLENEVARLKKLVGER |
| Swap3 | (GS) FSDTEEDDDEVIYSTKESMWDNLFPSDDALTIEAVKSLTPNNSTGSGSGSGSPESSDPAALK<br>RARNTEAARRSRARKLQRMKQLEDKVEELLCKNYHLENEVARLKKLVGER |
| Swap2 | (GS) ELSESFDSDDEPMINDSSAWTELFTNSDAPIEYNTVLKDTVDKTGSGSGSGSPESSDPAALK<br>RARNTEAARRSRARKLQRMKQLEDKVEELLCKNYHLENEVARLKKLVGER |
| Swap1 | (GS) EDVMDATYESSPIEPLSDVWKTLFDTFNDLASETSINNDTSKDEGSGSGSGSPESSDPAALK<br>RARNTEAARRSRARKLQRMKQLEDKVEELLCKNYHLENEVARLKKLVGER |
| 9acidA | (GS) STDSTPMFAYANLAANSKAWTSLFANAIPVTTAAVSLADKAIESGSGSGSGSPESSDPAALK<br>RARNTEAARRSRARKLQRMKQLEDKVEELLCKNYHLENEVARLKKLVGER |
| P+ | (GS) STDSTPMFEYENLEDNSKEWTPLFDNDIPVFTDDVSLADKPIDSGSGSGSGSPESSDPAALK<br>RARNTEAARRSRARKLQRMKQLEDKVEELLCKNYHLENEVARLKKLVGER |
| Aro2+ | (GS) STDSTPMFEYENLEDFEKNWTSLFDNDIQVTYDDVSLADKAIESGSGSGSGSPESSDPAALK<br>RARNTEAARRSRARKLQRMKQLEDKVEELLCKNYHLENEVARLKKLVGER |
| Aro3+ | (GS) STDYTPMFEYETYEDNFKEWTSLFDNDIPVTTDDVSLADRAIESGSGSGSGSPESSDPAALK<br>RARNTEAARRSRARKLQRMKQLEDKVEELLCKNYHLENEVARLKKLVGER |
| L123A+ | (GS) STDSTPMFEYENLEDNSKEWTSAFDNDIPVTTDDVSLADKAIESGSGSGSGSPESSDPAALK<br>RARNTEAARRSRARKLQRMKQLEDKVEELLCKNYHLENEVARLKKLVGER |
| L113A+ | (GS) STDSTPMFEYENAYDNSKEWTSLFDHDEPVTTDDVSLADKAIESGSGSGSGSPESSDPAALK<br>RARNTEAARRSRARKLQRMKQLEDKVEELLCKNYHLENEVARLKKLVGER |
| F108A+ | (GS) STDSTPMAEYENLEDNSKEWLSLFDNDIPVTTDDVSLADKPIESGSGSGSGSPESSDPAALK<br>RARNTEAARRSRARKLQRMKQLEDKVEELLCKNYHLENEVARLKKLVGER |
| W120A | (GS) STDSTPMFEYENLEDNSKEATSLFDNDIPVTTDDVSLADKAIESGSGSGSGSPESSDPAALK<br>RARNTEAARRSRARKLQRMKQLEDKVEELLCKNYHLENEVARLKKLVGER |
| R2 | (GS) STDSTPMFEYENLEDNSREWTSLFDNDIPVTTDDVSLADRAIESGSGSGSGSPESSDPAALK<br>RARNTEAARRSRARKLQRMKQLEDKVEELLCKNYHLENEVARLKKLVGER |
| WW+ | (GS) STDSTPMFNYENLEDNDKEWWSLFDWDIPVTWDDVSLADKAIESGSGSGSGSPESSDPAALK<br>RARNTEAARRSRARKLQRMKQLEDKVEELLCKNYHLENEVARLKKLVGER |
| solvvol | (GS) TTETTPMFEYENLEEQTKEWTTLFDQEIPVTTEEVTLADKAIETGSGSGSGSPESSDPAALK<br>RARNTEAARRSRARKLQRMKQLEDKVEELLCKNYHLENEVARLKKLVGER |
| SATtoG KtoR | (GS) GGDGGPMFEYENLEDNGREWGGLFDNDIPVGGDDVGLGDRGIEGGSGSGSGSPESSDPAALK<br>RARNTEAARRSRARKLQRMKQLEDKVEELLCKNYHLENEVARLKKLVGER |

| | |
|---|---|
| R3 | (GS) STRSTPSFEYENLEDNSKEWTSLFDNDIPVTTRDVSLADKAIRSGSGSGSGSPESSDPAALK<br>RARNTEAARRSRARKLQRMKQLEDKVEELLCKNYHLENEVARLKKLVGER |
| K5 | (GS) STKSTPMFEYENLEKNSKEWTSLFDNKIPVTTKDVSLADKAIKSGSGSGSGSPESSDPAALK<br>RARNTEAARRSRARKLQRMKQLEDKVEELLCKNYHLENEVARLKKLVGER |
| E3 | (GS) STDSEPMFEYENLEDNSEEWTSLFDNDIPVTTDDVSLADEAIESGSGSGSGSPESSDPAALK<br>RARNTEAARRSRARKLQRMKQLEDKVEELLCKNYHLENEVARLKKLVGER |
| ILVtoED | (GS) STDSTPMFEYENLEDNSKEWTSDFDNDEPETTDDESDADKADESGSGSGSGSPESSDPAALK<br>RARNTEAARRSRARKLQRMKQLEDKVEELLCKNYHLENEVARLKKLVGER |
| ILVtoED solvvol | (GS) TTETTPMFEYENLEEQTKEWTTDFDQEEPETTEEETDADKADETGSGSGSGSPESSDPAALK<br>RARNTEAARRSRARKLQRMKQLEDKVEELLCKNYHLENEVARLKKLVGER |
| ILVtoAS | (GS) STDSTPMFEYENLEDNSKEWTSSFDNDSPATTDDASSADKASESGSGSGSGSPESSDPAALK<br>RARNTEAARRSRARKLQRMKQLEDKVEELLCKNYHLENEVARLKKLVGER |
| StoL | (GS) STDLTPMFEYENLEDNLKEWTSLFDNDIPVTTDDVLLADKAIELGSGSGSGSPESSDPAALK<br>RARNTEAARRSRARKLQRMKQLEDKVEELLCKNYHLENEVARLKKLVGER |
| S/ArotoL | (GS) LTDLTPMLELENLEDNLKEWTSLFDNDIPVTTDDVLLADKAIELGSGSGSGSPESSDPAALK<br>RARNTEAARRSRARKLQRMKQLEDKVEELLCKNYHLENEVARLKKLVGER |
| E2K2 | (GS) STDKTPMFEYENLEDNEKEWTSLFDNDIPVTTDDVELADKAIEKGSGSGSGSPESSDPAALK<br>RARNTEAARRSRARKLQRMKQLEDKVEELLCKNYHLENEVARLKKLVGER |
| WTSLFD | (GS) STDSTPMWTSLFDEDNSKEWTSLFDNDIPVTTDDVSLADKAIESGSGSGSGSPESSDPAALK<br>RARNTEAARRSRARKLQRMKQLEDKVEELLCKNYHLENEVARLKKLVGER |
| ILVtoED W | (GS) STDSTPMFEYENLEDNSKEWTSLFDWDEPETTDDESDADKADESGSGSGSGSPESSDPAALK<br>RARNTEAARRSRARKLQRMKQLEDKVEELLCKNYHLENEVARLKKLVGER |
| R3 W | (GS) STRSTPSFEYENLEDNSKEWTSLFDWDIPVTTRDVSLADKAIRSGSGSGSGSPESSDPAALK<br>RARNTEAARRSRARKLQRMKQLEDKVEELLCKNYHLENEVARLKKLVGER |
| solvvol W | (GS) TTETTPMFEYENLEEQTKEWTTLFDWEIPVTTEEVTLADKAIETGSGSGSGSPESSDPAALK<br>RARNTEAARRSRARKLQRMKQLEDKVEELLCKNYHLENEVARLKKLVGER |
| $Gcn4^{cTAD30}$ | FEYENLEDNSKEWTSLFDNDIPVTTDDVSLGSGSGSGSPESSDPAALKRARNTEAARRSRARKLQR<br>MKQLEDKVEELLCKNYHLENEVARLKKLVGER |

FL Gcn4 (1-281):
GSMSEYQPSLFALNPMGFSPLDGSKSTNENVSASTSTAKPMVGQLIFDKFIKTEEDPIIKQDTPSNLDFDFALPQTATAPDAKTVL
PIPELDDAVVESFFSSSTDSTPMFEYENLEDNSKEWTSLFDNDIPVTTDDVSLADKAIESTEEVSLVPSNLEVSTTSFLPTPVLED
AKLTQTRKVKKPNSVVKKSHHVGKDDESRLDHLGVVAYNRKQRSIPLSPIVPESSDPAALKRARNTEAARRSRARKLQRMKQLEDK
VEELLCKNYHLENEVARLKKLVGER

$Med15^{7-659}$:
GSQDKDTLSNAERAKNVNGLLQVLMDINTLNGGSSDTADKIRIHAKNFEAALFAKSSSKKEYMDSMNEKVAVMRNTYNTRKNAVTA
AAANNNIKPVEQHHINNLKNSGNSANNMNVNMNLNPQMFLNQQAQARQQVAQQLRNQQQQQQQQQQQQQRRQLTPQQQQLVNQMKVA
PIPKQLLQRIPNIPPNINTWQQVTALAQQKLLTPQDMEAAKEVYKIHQQLLFKARLQQQQAQAQAQANNNNNGLPQNGNINNNINI
PQQQQMQPPNSSANNNPLQQQSSQNTVPNVLNQINQIFSPEEQRSLLQEAIETCKNFEKTQLGSTMTEPVKQSFIRKYINQKALRK
IQALRDVKNNNNANNNGSNLQRAQNVPMNIIQQQQQNTNNNDTIATSATPNAAAFSQQQNASSKLYQMQQQQQAQAQAQAQAQAQ
AQAQAQAQAAQAAQAQAQAQAQAQAQAQAQAQAQAQAQAQAQAQAHAQHQPSQQPQQAQQQPNPLHGLTPTAKDVEVIKQLSLD
ASKTNLRLTDVTNSLSNEEKEKIKMKLKQGQKLFVQVSNFAPQVYIITKNENFLKEVFQLRIFVKEILEKCAEGIFVVKLDTVDRL
IIKYQKYWESMRIQILRRQAILRQQQQMANNNGNPGTTSTGNNNNIATQQNMQ

*Protein expression and purification in E. coli*

All Gcn4 variants were expressed in *E. coli* BL21-Gold (DE3) or BL21(DE3) pLysS strains in ZYM5052 auto induction media [3] at 37°C for 24 hours. Cell pellets were resuspended in 1 M NaCl (pH 7.8), 30 mM imidazole and 10 mM 2-mercaptoethanol and lysed via sonication. Cell lysates were centrifuged, and the variants were purified from inclusion bodies as previously described [4]. The inclusion bodies were resolubilized in 6 M GdmHCl, 20 mM Tris (pH 7.5), 50 mM imidazole, 5 mM 2-mercaptoethanol overnight at 4°C. Solutions of solubilized inclusion bodies were cleared by centrifugation, and supernatants were loaded onto self-packed columns of chelating Sepharose fast flow beads (GE Healthcare) charged with nickel sulfate. The columns were washed with 4 column volumes of 4 M urea, 20 mM Tris pH 7.5, 50 mM imidazole and 1 mM DTT. Proteins were eluted from the Ni-NTA resin with 4 M urea, 20 mM Tris (pH 7.5), 500 mM imidazole, 1 mM DTT. TEV cleavage of the 6xHis-tag was done in 20 mM Tris (pH 7.5), 2 M urea, 50 mM

NaCl, 0.5 mM EDTA, 2 mM 2-mercaptoethanol overnight at 4°C. Cleaved protein solutions were loaded onto Ni-NTA columns. The flow-through and wash fractions were collected and concentrated using a 3k MWCO Amicon centrifugal filter. As a final purification step, the samples were passed in 2 M GdmHCl pH 7.3, 20 mM HEPES and 1 mM TCEP over a S75 Superdex size exclusion column (GE Healthcare). The identity of each protein was confirmed via intact mass spectrometry. All Gcn4 variants were stored in 20 mM HEPES (pH 7.3), 6 M GdmHCl, and 1 mM TCEP at 4°C.

$Med15^{7\text{-}659}$ was expressed in *E. coli* BL21-Gold (DE3) strain in LB media at 37°C, induced at $OD_{600}$ 0.8 with 1 mM IPTG, and the temperature was lowered to 18°C and cells were harvested the next day. Cell pellets were resuspended in 1 M NaCl (pH 7.8), 30 mM imidazole, 10 mM 2-mercaptoethanol and lysed via sonication. Cell lysates were centrifuged, and the supernatant was loaded onto self-packed columns as described above. The column was washed with 4 column volumes of 300 mM NaCl (pH 7.5), 50 mM imidazole, and 5 mM 2-mercaptoethanol. The protein was eluted with 500 mM NaCl (pH 7.5), 500 mM imidazole, and 5 mM 2-mercaptoethanol. TEV cleavage of the 6xHis-tag was done in 20 mM Tris (pH 7.5), 50 mM NaCl, 0.5 mM EDTA, and 1 mM DTT overnight at 4°C. Cleaved protein solutions were loaded onto Ni-NTA columns. The flow-through and wash fractions were collected and concentrated using a 3k MWCO Amicon centrifugal filter. To remove potential nucleic acid contamination, the protein solution was passed over a Heparin HP column (GE Healthcare). The column was pre-equilibrated with 20 mM Tris (pH 7.5), 50 mM NaCl, 5 mM 2-mercaptoethanol and the protein eluted with a linear salt gradient ending at 20 mM Tris (pH 7.5), 1 M NaCl, 5 mM 2-mercaptoethanol. As a final purification step, the protein solution was passed over a S200 Superdex size exclusion column (GE Healthcare) in 20 mM HEPES (pH 7.3) and 1 mM TCEP. The protein solution was stored at -80°C.

Given that Gcn4 variants were stored in denaturing buffer, we performed a rapid buffer exchange into 20 mM HEPES (pH 7.3), 2 mM DTT using 2 ml Zeba Spin desalting columns 7K MWCO (ThermoFisher) in preparation for all assays.

**Table S2: DNA constructs used in this study.**

| Name | Length (bp) | Sequence |
|---|---|---|
| 1HP | 12 | A**TGACTCA**TCGCGCAGCGATGAGTCAT |
| 2HP | 23 | A**TGACTCA**TATA**TGACTCA**TCGCGCAGCGATGAGTCATATATGAGTCAT |
| 4GRE | 41 | A**TGACTCA**TATA**TGACTCA**TTA**TGACTCA**TATA**TGACTCA**T |
| 8GRE | 83 | A**TGACTCA**TATA**TGACTCA**TTA**TGACTCA**TATA**TGACTCA**TTA**TGACTCA**TATA**TGACTCA**TTA**TGACTCA**TATA**TGACTCA**T |
| 1GRE | 41 | AGTACTCGTATACATCA**TGACTCA**TCATCTATAGCACTCGT |
| 2GRE | 41 | AGTACTCGTATA**TGACTCA**TTA**TGACTCA**TATAGCACTCGT |

Gcn4 response elements (GREs) are highlighted in green and hairpins in blue. For all sequences without hairpins, a reverse complement sequence was used to generate double-stranded DNA.

#### *Determination of saturation concentrations ($c_{sat}$) using HPLC*

The $c_{sat}$ values of all Gcn4 variants were determined via High-Performance Liquid Chromatography (HPLC) (Waters 2489 UV/Vis Detector) using the 280 nm detection channel. The input concentration to determine the homotypic saturation concentration ($c_{sat}$) of the Gcn4 variants was 200 µM. 60 µl of the protein solution was mixed with 20 µl of the phase separation buffer for a final buffer composition of 20 mM HEPES (pH 7.3), 150 mM potassium acetate, 10% PEG-8k, and 2 mM DTT. The samples were incubated at 20°C for 20 min, the protein solution was spun down (5 min, 12000 rpm, 20°C) and the dilute phase was transferred into a new tube. All samples were run over an analytical C4 HPLC column (Vydac). We equilibrated the column in $H_2O$ + 0.1% TFA. For elution, a linear gradient of acetonitrile (20-80%) was used. Additionally, we measured a standard curve of known protein concentration for the Gcn4 variants with different extinction coefficients.

Standard curve: $area_{\text{sample}} = m \times V_{\text{injected volume}} \times c_{\text{sat}} + b$

We determined the area under the curve using the built-in software on the HPLC. We then used the standard curve equation to determine the saturation concentrations. For more information on how to use the analytical HPLC method to quantify the amount of a biomolecule in a mixture in particular determining $c_{\text{sat}}$, please see [5]. At least three replicates per sample were measured and averaged.

*Fluorescence labeling*
DNA oligos which carried an Alexa-Fluor-488 label at the 5'-end were purchased (IDT sequence). Med15$^{7\text{-}659}$ was labeled with either LD-555 or LD-655 (maleimide) (Lumidyne Technologies) and Gcn4$^{\text{cTAD}}$ was labeled with Alexa Fluor 594 NHS (Succinimidyl Ester) (ThermoFisher).

*Microscopy analysis for in vitro phase separation*
All samples of Gcn4$^{\text{cTAD}}$ variants were prepared in 20 mM HEPES (pH 7.3), 150 mM potassium acetate, 10% PEG 8K, and 2 mM DTT. Protein concentrations were slightly above their corresponding $c_{\text{sat}}$ at 20 °C (in Fig. 2D). 2 μL of the protein solution was sandwiched between two coverslips with 3M 300 LSE high-temperature double-sided tape (0.34 mm) with a window for microscopy cut out. The samples were incubated at room temperature for 2 hrs to allow settling of condensates. Differential interference contrast (DIC) microscopy images were obtained at room temperature using a Nikon Eclipse Ni Widefield microscope with a 20x 0.75NA DIC N2 objective. For the DNA titration series, the final Gcn4$^{\text{cTAD}}$ concentration was 50 μM if not indicated otherwise. Due to the high viscosity of our PEG stock solution we pipetted PEG using a positive displacement pipette. All samples were measured in 20 mM HEPES (pH 7.3), 150 mM potassium acetate, 10% PEG 8K, 2 mM DTT and incubated at room temperature for 24 h to allow settling of condensates before taking DIC microscopy images. We determined the standard deviation of the pixel intensity using Fiji (Version 2.1.0/1.53o) as a measure of whether samples contained condensates. The standard deviation in the absence of condensates was 2.5 and is indicated by a dashed line in the plots.

*Confocal fluorescence microscopy*
To determine the fraction of labeled Med15$^{7\text{-}659}$ in Gcn4$^{\text{cTAD}}$ condensates in the absence and presence of DNA we used the same setup as for DIC, sandwiching the samples between double-sided tape (see above). The final concentrations were 50 μM Gcn4$^{\text{cTAD}}$ variant, 10 μM Med15$^{7\text{-}659}$ and 0.5 μM DNA$^{\text{4GRE}}$. The buffer conditions were 20 mM HEPES (pH 7.3), 150 mM potassium acetate, and 2 mM DTT. The samples were incubated for 24 h to allow settling of the condensates before imaging. Z-stack images were collected using a Zeiss LSM 980 Airyscan 2. All image analyses to determine the partition coefficient (PC) and dense phase volume fraction were performed using Fiji (Version 2.1.0/1.53o). The partition coefficient is defined as the ratio of the concentrations in the dense vs the dilute phase and calculated from the fluorescence intensities in the two phases, i.e., PC = $\text{intensity}_{\text{dense phase}}/\ \text{intensity}_{\text{dilute phase}}$. We created a mask of the dense phase based on fluorescence, determined the volume and average intensity of the dense phase. Some condensates had a very high PC (e.g., for 9acidA) or covered almost the entirety of the coverslip surface area, and there was only a small fraction of dilute phase. This can result in an under- and/or overestimation of $\text{intensity}_{\text{dilute phase}}$. To correctly determine $\text{intensity}_{\text{dilute phase}}$, we captured a separate plane that was 10 μm above the droplets. We determined the dense phase volume fraction by analyzing all slices of the z-stack that contained condensates, adding the

volumes, and dividing the sum by the total sample volume. The fraction of Med15$^{7\text{-}659}$ recruited into condensates is the product of the dense phase volume fraction and PC.

*Electrophoretic mobility shift assay (EMSA)*

Gcn4 protein variants to be tested for DNA binding affinity were adjusted to the following buffer conditions: 20 mM HEPES (pH 7.3), 100 mM KCl, 1 mM EDTA, 0.1% (v/v) NP40, 2 mM DTT and 10% (v/v) glycerol. Proteins were then flash frozen and stored in single use aliquots. Gcn4 variants were equilibrated at room temperature with Alexa Fluor 488–labeled 1GRE oligonucleotide probe at 5 nM final concentration in a volume of 10 μl. Electrophoresis was carried out in Mini-PROTEAN® Tetra cells (BioRad) using 1 mm 5% (w/v) polyacrylamide gels (acrylamide:bisacrylamide 37.5:1) in 0.5x TAE (20 mM Tris base, 10 mM glacial acetic acid and 0.5 mM EDTA). Gels were run at 100 V for 45 min on ice. After electrophoresis, fluorescence imaging of these gels was done on a ChemiDoc™ MP imager (BioRad).

*$K_D$ determination from fluorescence anisotropy measurements*

The dissociation constants of each of the Gcn4 protein variants to DNA were determined measuring changes in fluorescence anisotropy of the Alexa-Fluor-488 labeled 1GRE oligonucleotide probe upon protein binding. Measurements were carried out on a CLARIOstar plate reader (BMG Labtech) utilizing the FP-FITC protocol in precision mode with 200 flashes per sample. Binding assays were set up in black, small volume, Hibase 384-well microplates (Greiner Bio One #789400) in a final assay volume of 20 μl. In order to directly compare the results from the EMSA experiments, we used the same buffer (20 mM HEPES (pH 7.3), 100 mM KCl, 1 mM EDTA, 0.1% (v/v) NP40, 2 mM DTT and 10% (v/v) glycerol) to determine the dissociation constants of protein to DNA. The fluorescent probe concentration of 1HP was set to 50 pM final concentration.

For the dissociation constants of Med15$^{7\text{-}659}$ to a preformed Gcn4/DNA complex we used the same buffer conditions as in our phase separation microscopy studies (20 mM HEPES (pH 7.3, 150 mM potassium acetate, 2 mM DTT). The final concentration of each Gcn4 construct was 200 nM and 20 nM fluorescently labeled 1HP-Alexa-488 (if not indicated otherwise). Fluorescently labeled 1HP was purchased from IDT. Binding of Med15$^{7\text{-}659}$ to the preformed Gcn4/DNA complex slows down the tumbling of the labeled species, and this is detected by an increase in fluorescence anisotropy. We added increasing concentrations of Med15$^{7\text{-}659}$ to the preformed Gcn4/DNA complex. Measurements were done in triplicate and the data was analyzed using OriginLab.

$K_D$ values were obtained by fitting the data to the following equation introduced by Roehrl et al. [6]:

$$F_b = \frac{(K_D + 1HP + Med15) - \sqrt{(K_D + 1HP + Med15)^2 - 4(1HP) \times (Med15)}}{2 \times 1HP}$$

$F_b$ is the fraction bound of the fluorescent species, *1HP* is the total concentration of fluorescently labeled 1HP, *Med15* is the total concentration of Med15.

Measurements for all protein concentrations were done in triplicate. We observed a second transition for some of the variants (which likely represent the formation of larger complexes), which was excluded from the fit. All $K_D$ values are averages from three replicates with standard errors of the mean (S.E.M).

*Yeast strains and plasmid constructs*

A Gcn4 knockout strain constructed in BY4741 (MATa, his3Δ1, leu2Δ0, met15Δ0, ura3Δ0) was purchased from Horizon Discoveries (#YSC6273-201927482). The constructs designed for expressing all Gcn4 variants in *E. coli* were modified to insert an ATG start codon at the 5'end of the DNA sequence encoding the Gcn4 protein by Q5 site-directed mutagenesis (New England

Biolabs Inc). In addition to the Gcn4 expression constructs without further modification, a series of vectors was constructed wherein a 3xHA tag was added to the C-terminus of the Gcn4 coding region. For imaging purposes, some of these 3xHA-tagged expression constructs were further modified by inserting a GFP-encoding DNA fragment in frame into a unique restriction site within the 3xHA tag coding region. After sequence verification of the correct insertions, the DNA fragments encoding Gcn4 variants were cloned into the yeast expression vector pAG415Gal-ccdB (Addgene #14145) using a Gateway LR reaction (ThermoFisher). This vector allows for expression of the Gcn4 variant proteins in yeast by induction with Galactose. After cloning and sequence verification, constructs for expression of all Gcn4 protein variants were transformed into the Gcn4 knockout strain.

*Expression of Gcn4 variants in yeast*

Yeast colonies carrying plasmids for expressing Gcn4 variant proteins were used to inoculate 5 ml of Dropout medium without leucine, supplemented with 2% (w/v) Raffinose and grown at 30°C for two nights. After measuring $A_{600}$ of the initial culture, an appropriate volume of the starter culture was added to 10 ml fresh Dropout medium without leucine but with raffinose to result in an initial cell density equivalent to $A_{600}$ of 0.5. After growing these cultures for 1hr at 30°C, Galactose was added to a final concentration of 2% (w/v) for induction of Gcn4 protein expression, and growth was continued for an additional 4 hours at 30°C.

The cultures were then split into two samples each, one 1.5 ml sample for RNA purification, the remainder for extraction of total protein (see section below). Yeast RNA was purified utilizing a Nucleospin RNA plus kit (Takara/Macherey-Nagel #740984). For RNA purification, cells were pelleted at 4°C for 1 minute at 13,000 rcf. After removing the supernatant, about 200–250 μl of Zirconium beads were added to each tube followed by 400 μl of LBP buffer (supplied with the kit). Yeast cells were then broken by two 30” cycles of bead beating at the highest speed setting. In between cycles, the tubes were placed on ice for two minutes. After bead beating, samples were centrifuged for one minute at 13,000 rcf, 350 μl of supernatant were transferred to a DNA removal column and all RNA samples were processed further according to the instructions supplied with the kit. RNA concentrations and purity were determined using a Nanodrop spectrophotometer (ThermoFisher).

*Protein quantification from yeast expression*

Cell lysate preparation for protein expression was adapted from [7]. The cultures were centrifuged at 4000 rpm for 5 min at 4°C, washed once with water, then centrifuged as before. Pelleted cells were transferred to 1.5 ml microcentrifuge tubes, flash frozen, and stored at –80°C until further processing. Next, cells were thawed on ice and resuspended in 200 µl lysis Buffer (50 mM HEPES (pH 7.5), 140 mM potassium acetate, 1 mM EDTA, 1% Triton X-100, 0.1% sodium deoxycholate) supplemented with 1X Halt™ Protease Inhibitor Cocktail (Thermo Scientific) and 1X Halt™ Phosphatase Inhibitor Cocktail (Thermo Scientific). An equal volume of 400 μm zirconium beads (OPS Diagnostics) was added, and cells were subjected to bead beating for two rounds of 15 min at 4°C, with 15 min on ice between. A 22-gauge needle was heated using a Bunsen burner, then used to puncture a hole in the bottom of the microcentrifuge tube. The microcentrifuge tube was placed in a 2.0 ml tube and centrifuged at 7000 rpm for 30 sec to collect the lysate. Pelleted material was resuspended in the lysate and 200 μl was transferred to a fresh 1.5 ml microcentrifuge tube. Samples were supplemented with 10X DNase I Reaction Buffer (50 mM HEPES (pH 7.5), 25 mM $MgCl_2$, 5 mM $CaCl_2$) and treated with 5U DNase I (Thermo Scientific) at 37°C for 10 min. Samples were centrifuged at 13,000 rcf for 10 min at 4°C to pellet debris, and supernatant was transferred to a fresh 1.5 ml microcentrifuge tube. Laemmli Buffer (Bio-Rad) was added to each sample and an equal volume of each was run on a 4-20% Mini-PROTEAN®TGX™ precast gel (Bio-Rad). Following transfer, blots were stained for total protein content for 10 min with 0.1% (w/v) Ponceau S (Sigma-Aldrich) in 5% Acetic Acid (v/v). Blots were probed with antibody targeting Gcn4 (1:2000, Novus Biologicals) overnight at 4°C, and subsequently with rabbit anti-mouse HRP (1:5000, Abcam).

Gcn4 construct expression was verified by comparison to purified protein constructs as described above. Blot analysis was conducted using LI-COR Image Studio™. Gcn4 levels were normalized to total protein content (20-60 kDa), as quantified by Ponceau S staining, then corrected for interfering non-specific bands also present in the vector strain. Final protein values were calculated as a fold change with respect to Gcn4$^{cTAD}$.

*RT-qPCR analysis*

Expression levels of GCN4-induced genes were analyzed by RT-qPCR. Five hundred nanograms of RNA from each sample were reverse transcribed using LunaScript RT Master mix in a final volume of 20 μl. Following reverse transcription, cDNA samples were diluted by addition of 230 μl of RNAse free water, and 2.5 μl of each diluted sample was then used per qPCR reaction. To each reaction, 2.5 μl of each primer pair (200 nM final concentration) and 5 μl 2x Luna Universal qPCR Master mix were added to a final volume of 10 μl. RT-qPCR was performed on a QuantStudio 6 instrument (Applied Biosystems) for 40 cycles in a 384-well format, and it included a melting curve analysis. Ct (Cq) values were determined with the Software package "Design and Analysis 2.5.1" which was supplied with the instrument, thresholds were only adjusted as necessary. Relative mRNA expression levels were then calculated by the comparative $C_T$ method [8]. The expression levels of ALG9, TAF10 and UBC6 are among the most stable under a wide variety of growth conditions in yeast [9] and their average Cq was used for normalization of the mRNA expression levels of GCN4 activated target genes.

**Table S3: The following primer pairs were used for the RT-qPCR analysis.**

| Primer | Sequence |
|---|---|
| ALG9-F | 5′-CCG TTG CCA TGT TGT TGT ATG-3′ |
| ALG9-R | 3′-GCC AGG AAA TTG TAC GCT AAA C-5′ |
| TAF10-F | 5′-GTC TTC CGT AGC GGT ATC TAA TG-3′ |
| TAF10-R | 3′-CTG TTG ATG TTG TTG TTG TGA AAT C-5′ |
| UBC6-F | 5′-CGG CAA ATA CAG GTG ATG AAA C-3′ |
| UBC6-R | 3′-TCA GCG CGT ATT CTG TCT TC-5′ |
| ARG1-F | 5′-GCT GGC AGA AAG GAT TTG TTA G-3′ |
| ARG1-R | 3′-CCA AGA TAC CTG CCT CGT AAG-5′ |
| ARG3-F | 5′-CGC ATG TCT GAA ATT CGG TAT AAG-3′ |
| ARG3-R | 3′-CAA ATG TCG CAC CGT TTC TC-5′ |
| ARG4-F | 5′-CCA TGA AGG TTT GGC TGA AAT C-3′ |
| ARG4-R | 3′-GAT CTA CCG GTG TGG ACT TTA C-5′ |
| HIS4-F | 5′-GCT GCC GAT TTG TTC TAC TTT G-3′ |
| HIS4-R | 3′-TAG CAT CAC CTT TCC GTC TTG-5′ |
| CPA2-F | 5′-GCT GCT GAA AGG GTC AAA TAC-3′ |
| CPA2-R | 3′-GCG GCA AGT TCC TTC ATT TC-5′ |
| MET16-F | 5′-TTG CAC CAT TTC CCA CAA AC-3′ |
| MET16-R | 3′-CCT CCG ATT CAC ATC CAT CC-5′ |

*Gcn4 target gene promoter analysis*

Gcn4 target gene promoters used in activity value calculations were analyzed for Gcn4 [10] and histone H3 [11] occupancies. Bigwig files for three replicates of Gcn4 and H3 in uninduced and SM induced conditions were aligned to the sacCer3 genome in IGV. Gcn4 motifs were identified using the Find Motif tool, with the Gcn4 binding motif (TGAsTCA) used as the input. One replicate of each was used as a representative in Figure S1.

*Yeast cell imaging*

Strains expressing the C-terminal GFP-tagged Gcn4 variants were inoculated in 5 mL of -LEU + 2% raffinose synthetic media and were grown overnight at 30°C with shaking. The next day, these pre-cultures were diluted in 10 mL fresh -LEU + 2% raffinose synthetic media to $OD_{600}$ ~ 0.3 and allowed to grow at 30°C with shaking for 6h. This was followed by induction for 2h with a final concentration of 2% Galactose at 30°C with shaking. After induction 1 mL culture was spun at 500 g at room temperature for 1 minute. The supernatant was discarded, and the obtained pellet was resuspended in 1 mL of fresh -LEU + 2% raffinose synthetic media. 25 μL of this cell suspension was spotted on the coverslip and allowed to settle for 5-10 minutes. The extra cell suspension was aspirated using a micropipette, and mounting was done with 5-10 μL of ProLong Diamond antifade mounting agent containing DAPI (Invitrogen, P36790). The coverslip was sealed with nail polish. Z-stack images were acquired on a Zeiss LSM 980 Airyscan 2 with *63x*/1.4 oil immersion objective, and images were further zoomed in to 6X. Image processing and analysis were done using FIJI (Version 2.14.0/1.54f). 200 cells from each variant were imaged and the ratio of nuclear-to-cytoplasmic localization of Gcn4 variants was calculated using the formula N/C= Total nuclear fluorescence intensity/Total cytoplasmic fluorescence intensity [12] and was plotted with standard error.

*Chromatin Immunoprecipitation followed by qPCR (ChIP-qPCR)*

Overnight cultures of each strain in 50 mL dropout medium without leucine, supplemented with 2% (w/v) Raffinose were diluted to $OD_{600}$ = 0.5 in identical media and grown for two hours with shaking (180 rpm) at 30°C. Galactose was added to a final concentration of 2% (w/v) and cells were incubated at 30°C with shaking for 2 hrs. Following induction, formaldehyde was added to 1% (v/v) final concentration, and the culture was incubated at RT for 15 min with constant shaking at 70 rpm. Formaldehyde was then quenched by adding glycine to a final concentration of 125 mM for 5 min at RT with shaking at 70 rpm. Cultures were transferred to 50 ml falcon tubes and centrifuged at 4000 rpm for 5 minutes at 4°C. Media was discarded and cells were resuspended in 1 ml cold water and transferred to a microcentrifuge tube. After centrifugation, supernatants were discarded, flash frozen in liquid nitrogen and stored at –80°C until processing.

For processing, cell pellets were thawed on ice and resuspended in 500 μl of Sonication Buffer (50 mM HEPES-KOH pH 7.5, 140 mM NaCl, 4 mM EDTA, 2 mM EGTA, 1% Triton X-100, 0.1% Sodium deoxycholate, 0.1% SDS). Next, 200 μl silica beads (OPS Diagnostics) were added, and cells were broken by two cycles of bead beating at 3000 rpm for 30 minutes in the cold room, interrupted by a cool down period of 15 minutes. The supernatant was isolated by piercing the tube bottom with a 22-gauge needle, placing in a 2 ml tube, and centrifuging at 7000 rpm for 30 s. The visible pellet was resuspended and the entire volume was transferred to a Covaris 1 ml microtube. Samples were sheared using a Covaris S220 sonicator (20% DF / 240w PIP / 200 CPB / 4°C / 2 min continuous shearing). Following sonication, the sample was transferred to a fresh 1.5 ml microcentrifuge tube and centrifuged for 10 min at 13,000 rpm at 4°C, after which the supernatant was transferred to a new 1.5 ml microcentrifuge tube (whole cell lysate). A 1% input sample was reserved at this stage and the remaining supernatant was used in the subsequent immunoprecipitation (IP). For each IP, 5 μl of anti-HA antibody (Cell Signaling #3724) was coupled to Epoxy M270 beads per manufacturer protocol (Invitrogen) and added to the whole cell lysate. After rotating samples overnight at 4°C, beads were washed on ice twice with 1 ml Sonication Buffer, once with 1 ml High Salt Sonication Buffer (50mM HEPES-KOH pH 7.5, 500 mM NaCl, 4mM EDTA, 2mM EGTA, 1% Triton X-100, 0.1% Sodium deoxycholate, 0.1% SDS), once with 1 ml LiCl Buffer (40 mM Tris-Cl pH 7.5, 2mM EDTA, 500 mM LiCl, 1% NP-40, 1% sodium deoxycholate), and once with 1 ml TE supplemented with 0.1% Triton X-100 (10 mM Tris pH 8.0, 1 mM EDTA, 0.1% Triton X-100). Beads were then resuspended in 200 μl Elution Buffer (50 mM

Tris pH 8.0, 1 mM EDTA, 1% SDS) and heated at 65°C for 30 min with shaking at 850 rpm. The supernatant was transferred to new 1.5 ml microcentrifuge tube. The input samples were thawed and diluted to 200 µl final volume with Elution buffer. Samples were then incubated overnight at 65°C without shaking. The following day, samples were treated with RNase A (Sigma) at 37°C for 2 hrs, followed by Proteinase K (Invitrogen) treatment at 55°C for 30 min and phenol-chloroform-isoamyl alcohol extraction. After ethanol precipitation, the extracted DNA was resuspended in 100 µl TE Buffer (10 mM Tris-HCl pH 8.0, 0.1 mM EDTA).

For qPCR analysis 25 µl of each resuspended DNA sample was diluted with 100 µl of water, and 2.5 µl of the diluted DNA was then used per reaction, and qPCR was performed as described above. The primer pairs indicated in Table S5 for each promoter region were used.

**Table S4: Primer pairs for ChIP-qPCR analysis.**

| Primer | Sequence |
|---|---|
| CPA2prom-F | 5′- CCCTATCAGAAATATTCAAATGTCC -3′ |
| CPA2prom-R | 3′- CTATAGAATGCCGCAGGAAAG -5′ |
| ARG1prom-F | 5′- GCCCAGCCACTGAGATAAG -3′ |
| ARG1prom-R | 3′- GTTAATGGAAATAGGTTGCGACAG -5′ |
| ARG3prom-F | 5′- TCGTATTACTCATTCAGCTCTTCC -3′ |
| ARG3prom-R | 3′- TTTGTCATCTCGAAACGAAACG -5′ |
| ARG4prom-F | 5′- TGGTTACTCATTGGCAGAATCC -3′ |
| ARG4prom-R | 3′- AGTTCTGTGCTTCGCTGAC -5′ |
| HIS4prom-F | 5′- GAACTGACTCTAATAGTGACTCC -3′ |
| HIS4prom-R | 3′- CACAGTATACTACTGTTCATAGTC -5′ |
| MET16prom-F | 5′- CATACTGTTCTTTATTCCGTCG -3′ |
| MET16prom-R | 3′- AGGTTACAAACAGTACAGTATAC -5′ |

### Supplementary Figures

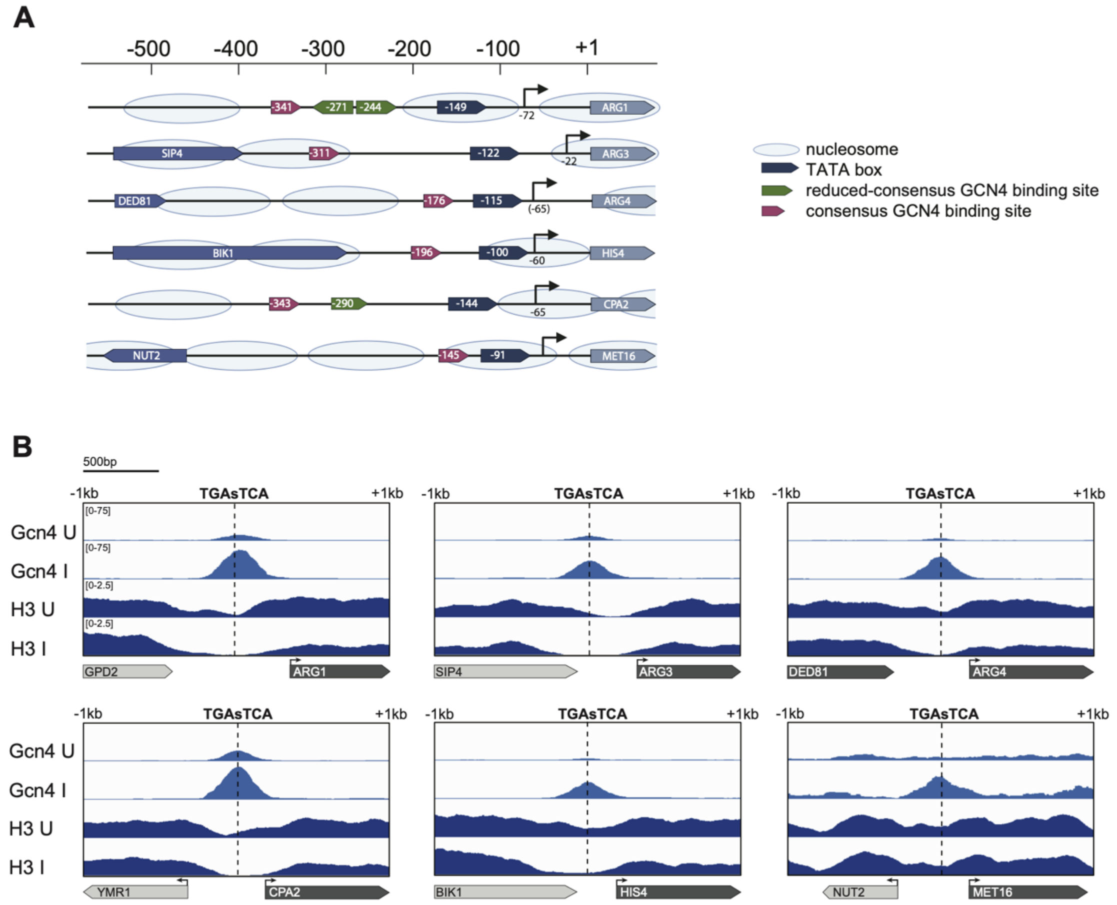


**Figure S1: Structure of promoter regions of Gcn4 target genes and their accessibility and occupancy. (A)** Structure of the promoter regions of the six Gcn4 target genes whose activation was monitored. Target genes are aligned at their translational start codons (+1). Locations of TATA boxes, transcriptional start sites (indicated by an arrow from SAGE analysis; position in bracket is extrapolated from primer extension) and GCN4 binding sites whose sequences provide a perfect match to the consensus binding site (RTGASTCAY) or a match to a reduced seven nucleotide binding site consensus (TGACTCA) are shown**.** Only occupied binding sites that are identified by ChIP-Seq analysis [10] are shown, although additional potential binding sites can be identified by sequence analysis in the promoter regions. The light blue ovals depict the approximate locations of nucleosomes in the uninduced state (data extracted from [13]). **(B)** ChIP-Seq traces produced with anti-GCN4 and anti-Histone H3 antibodies from yeast cells in the uninduced state (U), induced state (I) and in GCN4 knockout cells at the GCN4 binding sites of the promoter regions for the six target genes from A from Reference [10]. Upon GCN4 induction, the traces show an increase in the occupancy of the GCN4 binding sites and a concurrent decrease in H3 occupancy resulting in creation or widening of a nucleosome-free region in most target promoter loci.

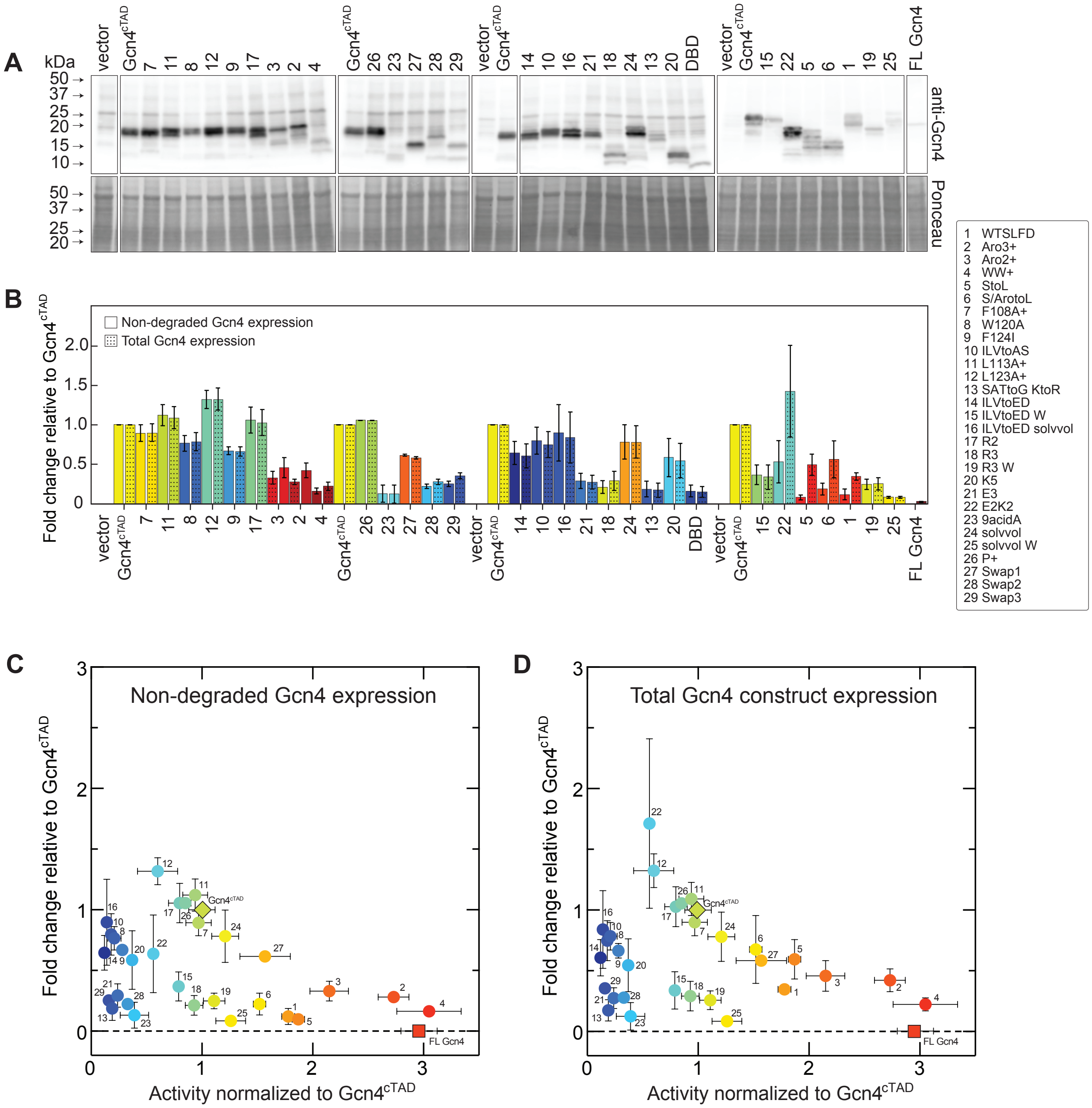


**Figure S2: Expression levels of GCN4 variants *in vivo* and correlation with target gene activation. (A)** Immunoblots of whole cell extracts from GCN4Δ yeast strains expressing the indicated Gcn4 construct. Blots were probed with an anti-Gcn4 antibody raised against the Gcn4 DNA-binding domain (top) and stained with Ponceau S (bottom) for total protein content. **(B)** To account for whether degradation of Gcn4 protein variants contributes to differences in target gene activation, we quantitated signals for full-length (non-degraded) and total Gcn4 protein levels for each construct, subtracted non-specific background taken from the vector only control lane and normalized it to total protein content. Numbers are presented as fold change relative to Gcn4$^{cTAD}$. The values are the mean of three independent samples ± SEM. Correlation plots between Gcn4 protein levels and target gene expression levels are shown for non-degraded Gcn4 protein expression **(C)** or total protein expression **(D)**.

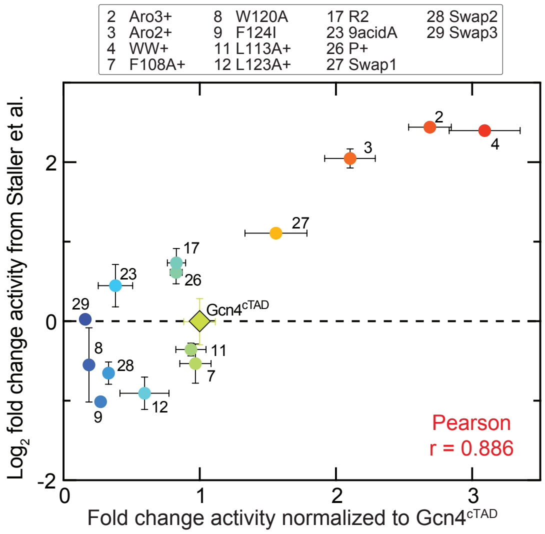


**Figure S3: Comparison of activities for a subset of Gcn4 variants measured in this study to activities reported previously [1].** The Pearson correlation coefficient was calculated to be 0.886.

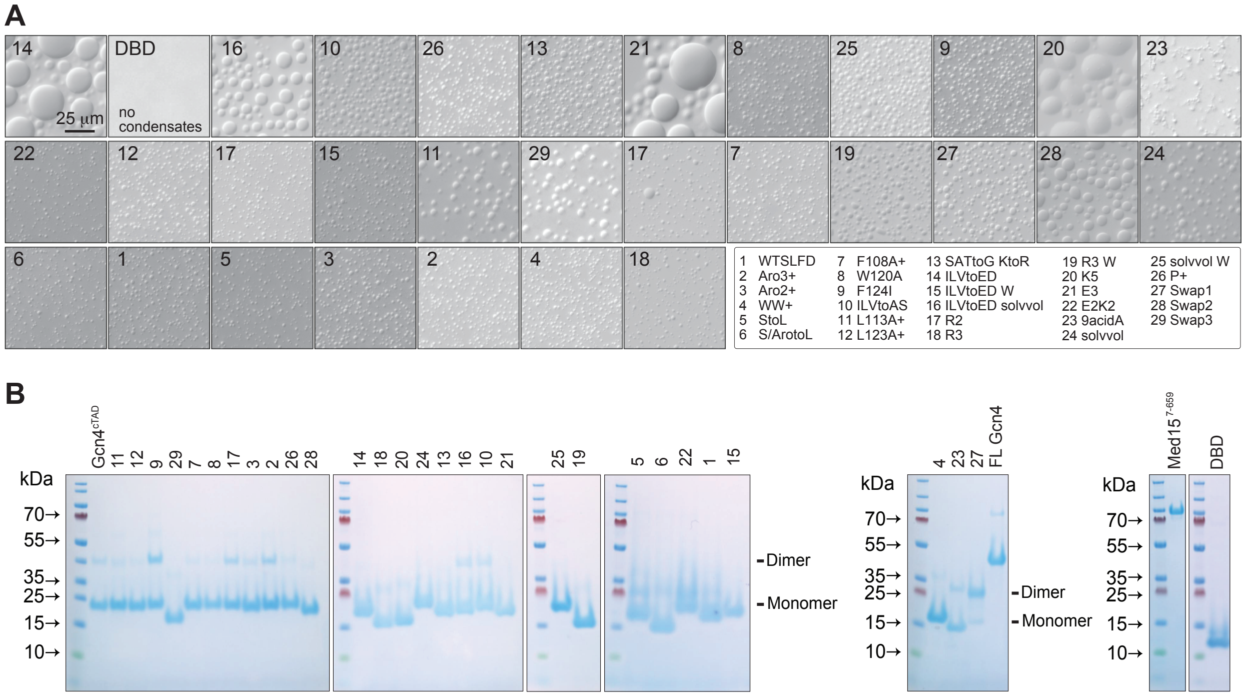


**Figure S4: Purified proteins used in this study. (A)** SDS-PAGE analysis of purified proteins. A second band at higher molecular weight (representing a dimer) was detected for some Gcn4 variants. **(B)** DIC images of Gcn4 variants showing that all variants can undergo phase separation and form condensates except DBD, which did not form condensates up to 0.8 mM. Solution conditions were 20 mM HEPES (pH 7.3), 150 mM potassium acetate, 10% PEG 8K, 2 mM DTT. The protein concentrations are not comparable; the images serve to indicate the presence of condensates.

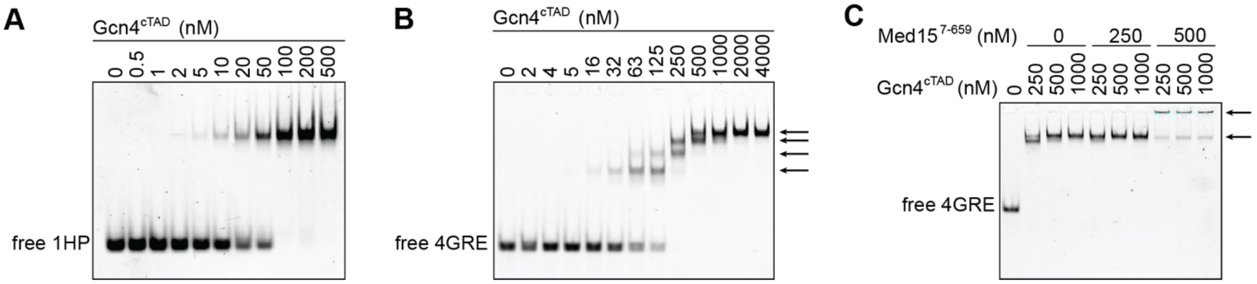


**Figure S5: EMSA of Gcn4$^{cTAD}$ binding to oligonucleotide probes shows the formation of higher-order complexes**. **(A)** A fluorescently labelled oligonucleotide probe containing one consensus Gcn4 binding site (1HP) was incubated with increasing concentrations of Gcn4$^{cTAD}$. **(B)** A fluorescently labelled oligonucleotide probe containing four consensus Gcn4 binding sites (4GRE) was incubated with increasing concentrations of Gcn4$^{cTAD}$. The arrows on the right point to the positions of four distinct protein–DNA complexes corresponding to species with one to all four binding sites filled. **(C)** The same probe used in panel B was incubated with Gcn4$^{cTAD}$ and Med15$^{7\text{-}659}$ at the concentrations shown. Upon addition of Med15$^{7\text{-}659}$, Gcn4 protein–DNA complexes change their mobility suggesting formation of high molecular weight complexes with Med15$^{7\text{-}659}$ indicated by arrows. Buffer conditions: 20 mM HEPES (pH 7.3), 100 mM KCl, 1 mM EDTA, 0.1% (v/v) NP40, 2 mM DTT and 10% (v/v) glycerol.

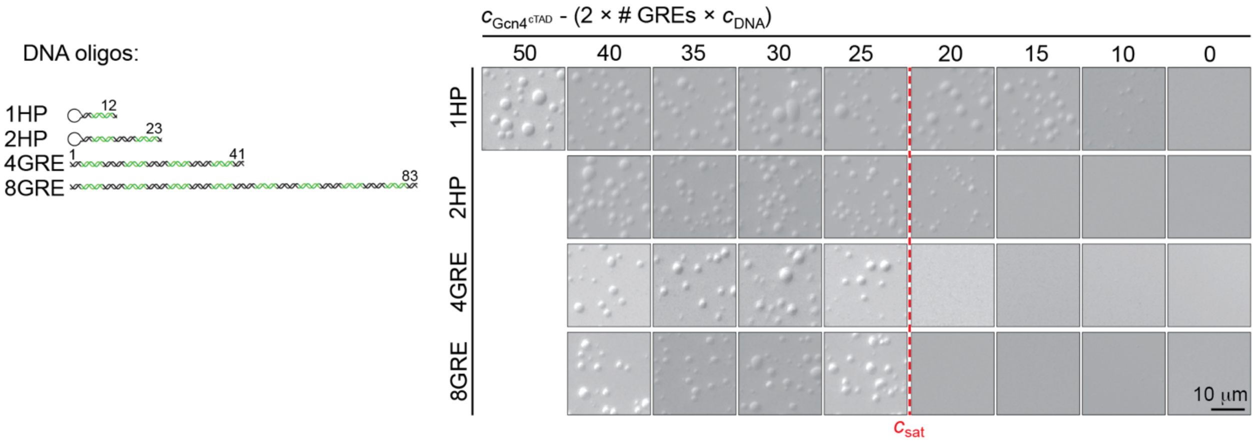


**Figure S6: Dissolution of Gcn4$^{cTAD}$ condensates with increasing concentration of DNA oligos of varying length.** DNA oligos varied in length and # of GREs from one up to eight GREs (indicated on the left). Gcn4 condensates disappear when the concentration of free Gcn4$^{cTAD}$ drops below its saturation concentration ($c_{sat}$) except for shorter DNA oligos. There the Gcn4 condensates dissolve at higher DNA concentration. The saturation concentration is indicated by the red dashed vertical line. Solution conditions were 20 mM HEPES (pH 7.3), 150 mM potassium acetate, 10% PEG 8K, 2 mM DTT.

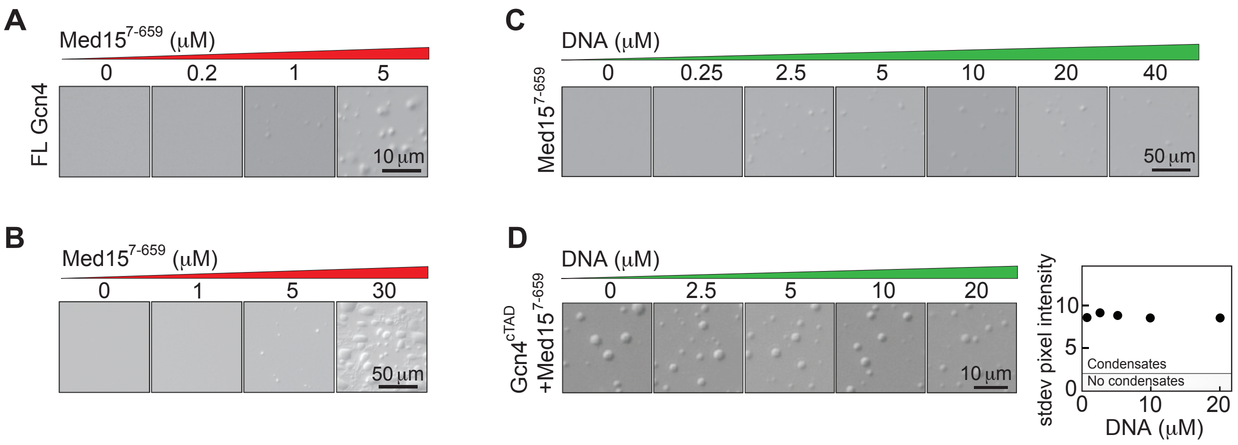


**Figure S7: Mediator subunit Med15 is important factor in mediating multicomponent phase separation in Gcn4 and DNA. (A)** Med15$^{7-659}$ enhances phase separation of Gcn4$^{FL}$ in the presence of crowders *in vitro*. **(B)** Med15$^{7-659}$ forms condensates in the presence of crowders by itself *in vitro*. **(C)** Med15$^{7-659}$ forms condensates with DNA$^{4GRE}$ oligonucleotide in the presence of crowders. **(D)** Titration of DNA$^{4GRE}$ oligonucleotide into Gcn4$^{cTAD}$/Med15$^{7-659}$ samples does not affect condensates. Index of dispersion remains the same with increasing 4GRE concentration. Solution conditions were 20 mM HEPES (pH 7.3), 150 mM potassium acetate, 2 mM DTT with or without 10% PEG 8k as indicated.

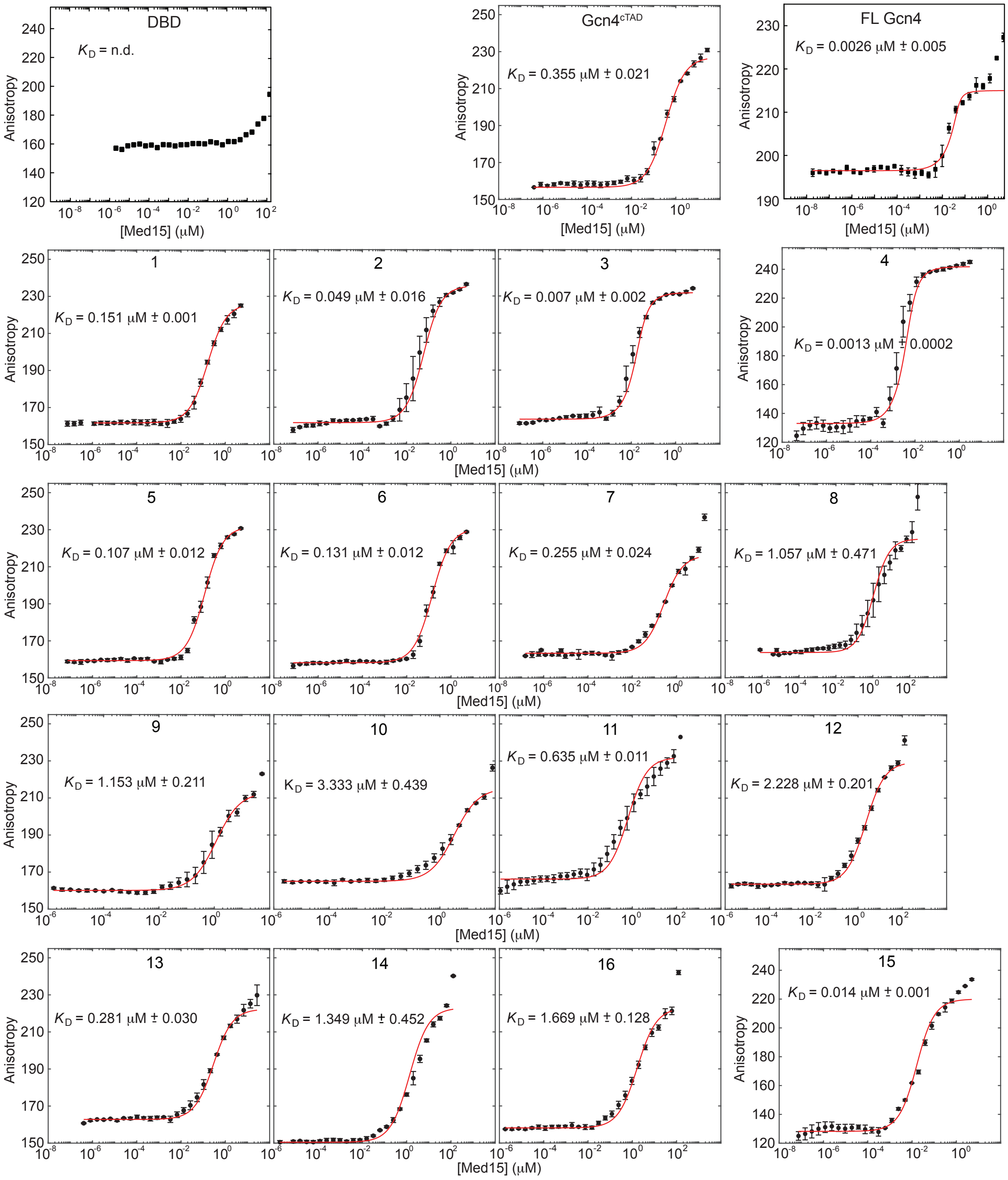

DBD
$K_D$ = n.d.
Gcn4cTAD
$K_D$ = 0.355 μM ± 0.021
FL Gcn4
$K_D$ = 0.0026 μM ± 0.005
1
$K_D$ = 0.151 μM ± 0.001
2
$K_D$ = 0.049 μM ± 0.016
3
$K_D$ = 0.007 μM ± 0.002
4
$K_D$ = 0.0013 μM ± 0.0002
5
$K_D$ = 0.107 μM ± 0.012
6
$K_D$ = 0.131 μM ± 0.012
7
$K_D$ = 0.255 μM ± 0.024
8
$K_D$ = 1.057 μM ± 0.471
9
$K_D$ = 1.153 μM ± 0.211
10
$K_D$ = 3.333 μM ± 0.439
11
$K_D$ = 0.635 μM ± 0.011
12
$K_D$ = 2.228 μM ± 0.201
13
$K_D$ = 0.281 μM ± 0.030
14
$K_D$ = 1.349 μM ± 0.452
16
$K_D$ = 1.669 μM ± 0.128
15
$K_D$ = 0.014 μM ± 0.001
Anisotropy
[Med15] (μM)

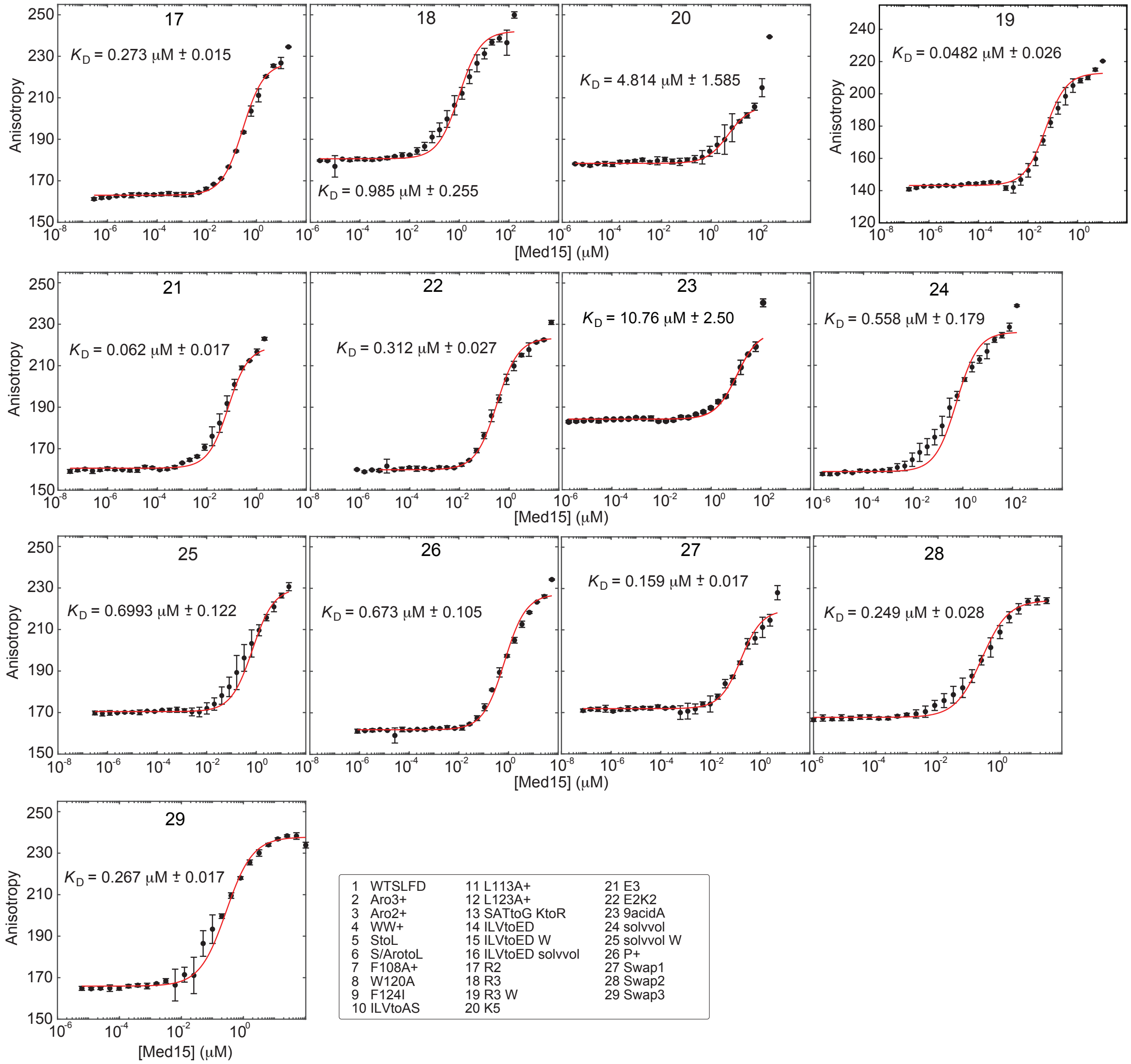


**Figure S8: Determination of dissociation constant ($K_D$) of Med15 to preformed Gcn4–DNA complex via fluorescence anisotropy.** Med15 was titrated into 1HP fluorescently labeled DNA (20 nM) and Gcn4 (200 nM). The dissociation constant was determined for the first transition using equation 5 from ([6]). The fit is shown as a red line. Error bars indicate ± S.D. from at least three independent measurements. Solution conditions were 20 mM HEPES (pH 7.3), 150 mM potassium acetate, 0.1% (v/v) NP40 and 2 mM DTT. No binding of Med15 to preformed DBD–DNA complex was detected up to 300 μM of Med15.

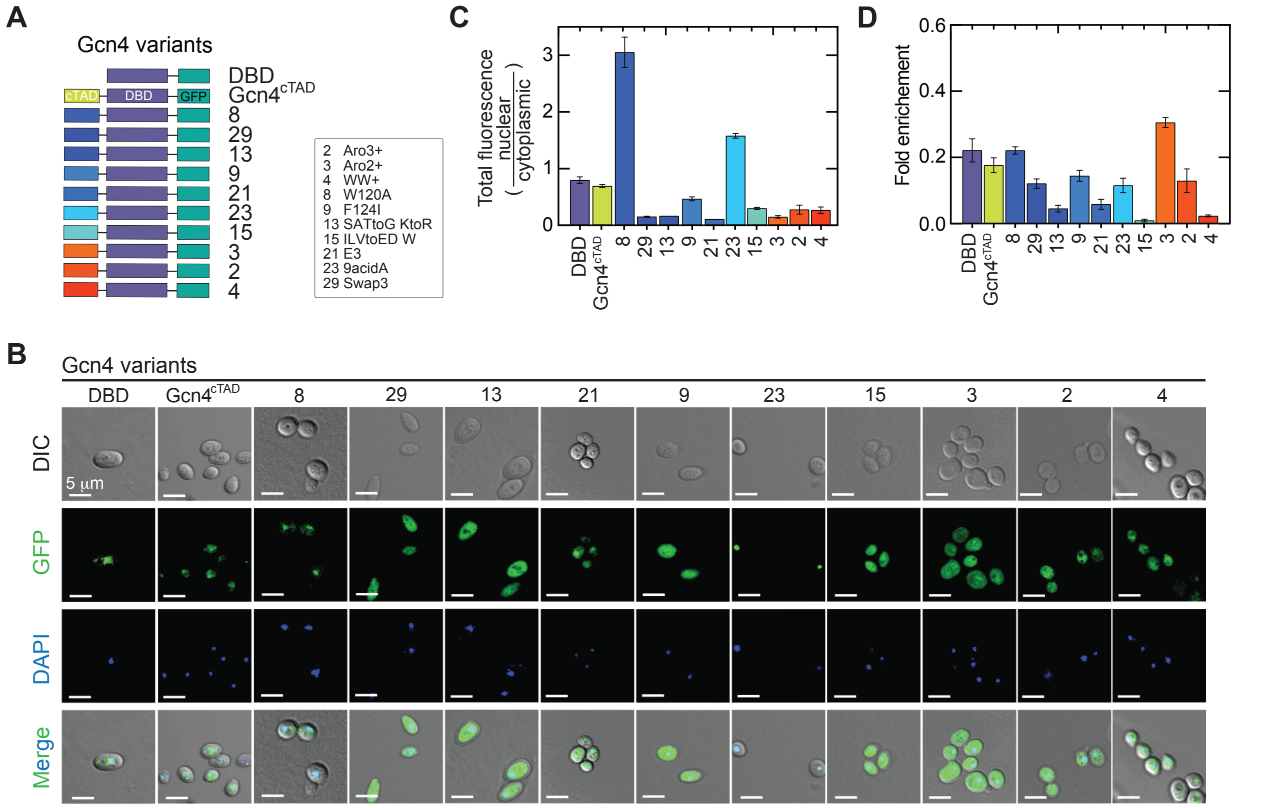


**Figure S9: Mislocalization of Gcn4 variants may contribute to low activity. (A)** Schematic structure of Gcn4 variants labelled with GFP at the C-terminus of the DNA binding domain and examined for nuclear localization *in vivo* by confocal microscopy. **(B)** Representative DIC and confocal fluorescence micrographs showing nuclear and cytoplasmic distribution of Gcn4 protein variants (green) in yeast cells. Nuclei were stained with DAPI (blue). **(C)** Quantitation of green-fluorescent signals to measure distribution ratio of Gcn4 variants in yeast cells. **(D)** ChIP analysis of target engagement by Gcn4 variants at target promoters. Each target was analyzed via three independent replicates, and fold enrichment of promoter fragments over genomic DNA input was calculated and averaged for the six target genes shown in Fig. 1C. Error bars in C and D indicate ± S.E.M. from at least three independent measurements.
